# From adolescence to adulthood: functional fingerprints of high-level visual cortex reveal differential development of visuospatial processing

**DOI:** 10.64898/2026.02.11.705468

**Authors:** Jewelia K. Yao, Justin Choo, Dawn Finzi, Kalanit Grill-Spector

## Abstract

Reading, social interaction, and navigation rely on visuospatial computations by population receptive fields (pRFs) in category-selective regions of the ventral, lateral, and dorsal streams. However, how visuospatial computations vary across streams and category, or develop during adolescence is unknown. Using functional magnetic resonance imaging (fMRI) and pRF modeling in adolescents and adults, we investigate the development of pRFs and category-selectivity. Adolescents and adults show a consistent functional fingerprint whereby, pRF location, pRF size, and visual field coverage vary by category-selectivity, stream, and hemisphere. While pRF location is largely adult-like by adolescence, pRF size, visual field coverage, and category-selectivity exhibit region-specific increases and decreases from adolescence to adulthood. Together, these findings delineate a timeline of continued functional plasticity during adolescence and provide a multidimensional framework for understanding the organization of high-level visual cortex.

## Introduction

Across development, the ability to perceive faces, words, bodies, and places is crucial for key behaviors like social interactions, reading, and navigation, and depends on computations by neurons throughout visual cortex^1–5^. In humans, perception of these categories is enabled by computations in category-selective regions across three distinct visual processing streams: the ventral stream, which extends ventrally from occipital to temporal cortex and is involved in visual recognition^1,2^, the dorsal stream, which runs through superior occipital-parietal cortex and is engaged in spatial navigation and attention^1,2^, and the lateral stream, which extends from lateral occipitotemporal cortex through the superior temporal sulcus (STS) and is involved in dynamic, action, and social perception^3–6^. While many studies examined functional differences among the three pathways, few have investigated how basic visual computations such as visuospatial processing by population receptive fields (pRFs) – the portion of visual space processed by the population of neurons in a voxel^7^– differ within and across streams. Even less is known about how pRFs in category-selective regions develop, particularly in adolescence when visual behaviors like face recognition and reading, and the visual areas supporting them, are still developing^8–10^. Thus, we ask: (1) Do pRFs in category-selective regions develop during adolescence? (2) How do pRFs differ across category-selective regions and streams?

In each visual area, pRFs are organized systematically, tiling the visual field; this is referred to as the visual field coverage (VFC) of an area^11^. While much is known about VFC and pRF properties in early retinotopic areas^12,13^, pRFs in category-selective regions in the ventral, lateral, and dorsal streams have been less studied, partly because traditional pRF mapping experiments used flickering checkerboards designed to activate early and intermediate, rather than high-level visual areas^7,14–18^. More recent studies in adults have employed pRF mapping stimuli that include shapes, objects, faces, and colors that drive neurons in high-level regions, enabling the estimation of pRFs in category-selective regions^11,19–24^. Findings of differential retinotopic biases and pRFs in high-level visual cortex led researchers to hypothesize about the nature of these differences.

With respect to category, researchers have hypothesized that retinotopic biases in category-selective regions reflect fixation patterns for respective categories. For example in adults’ ventral face- and word-selective regions, there is a higher concentration of pRFs and VFC around the center of gaze (fovea)^20,25–27^, paralleling adults’ looking behavior, which involves fixating on faces and words in order to recognize faces and read. Further, when people fixate on faces, bodies will be below the face, and this is mirrored in the lower-visual field bias of pRFs of ventral body-selective regions^28^. Additionally, in adults’ ventral place-selective areas, pRFs and VFC extend into the periphery^20,26,27,29,30^, mirroring the real world in which places encompass the entire visual field. The category hypothesis thus suggests that pRF differences across category-selective regions are driven by unique spatial regularities and distinct viewing patterns that are associated with different categories^22,26,27,31,32^. With respect to streams, researchers have hypothesized that different streams have distinct computational objectives^1–5^ that utilize visual information from different parts of the visual field.

Thus, the stream hypothesis predicts that pRFs will systematically vary across streams. Some studies suggest differences in upper/lower visual field biases across streams with ventral stream pRFs concentrated in the upper visual field and dorsal stream pRFs concentrated in the lower visual field^33,34^. Other studies suggest differences in eccentricity biases across streams, whereby pRFs and VFC are concentrated around the fovea in ventral face-selective regions but extend to the periphery in lateral face-selective regions^3,20^. Nonetheless, the category and stream hypotheses are not mutually exclusive, as pRFs and VFC may vary by both category and stream.

The category and stream hypotheses offer frameworks for how pRF properties and VFC might differ across visual cortex in adults, but they do not make predictions regarding developmental trajectories. Adolescence, the period between ages 10 to 19 years, presents a unique developmental window as crucial visual behaviors, like face recognition, reading, and spatial attention, along with the underlying category selectivity in face-, limb-, and word-regions, are still developing^9,35–38^. Currently, we have no knowledge of the development of visuospatial processing after age 12^25,39–41^. Thus, understanding how pRFs may develop during adolescence will provide important insights into the developmental timeline of basic visual functions.

We consider two possibilities regarding pRF development: one possibility is that as category selectivity develops into adolescence^9,35–37^, pRF properties in high-level visual areas will also continue to develop. This hypothesis predicts that pRF properties and VFC in category-selective regions of adolescents will differ from that of adults and is supported by studies finding that pRFs in category-selective regions of VTC, pFus-faces and pOTS-words, continue to develop from age 5 to adulthood^25^. Alternatively, as pRFs perform basic spatial computations on visual inputs, they may mature before higher-level, category computations, and be fully developed by adolescence. This hypothesis predicts no significant differences between adolescents and adults in pRFs and VFC in category-selective regions, and is supported by work demonstrating that pRF properties and VFC of early visual areas (V1-V3) are adultlike as early as 5 to 7 years of age^25,39,40^. Of course, it is also possible that pRFs develop differentially across regions, with pRFs and VFC developing in only some regions during adolescence.

## Results

### Toonotopy drives high-level category-selective regions in adolescents and adults

To test these hypotheses, 15 adolescents (ages 10-17; 9 females, 6 males) and 27 adults (ages 22 - 32; 13 females, 14 males) participated in two fMRI experiments: one to map pRFs using sweeping bars with cartoons (Toonotopy^20^; **Fig. 1A**) and another to identify category-selective regions (functional localizer^42^; **Supplementary Figure 1A**). There were no significant differences between adolescents and adults in motion (all subjects moved < 2 voxels) or task performance accuracy during the Toonotopy experiment (adolescents: M = 97 ± 5.6, adults: M = 93.14 ± 9.2; t(8.48) = 1.01, p = 0.34).

**Figure 1.**
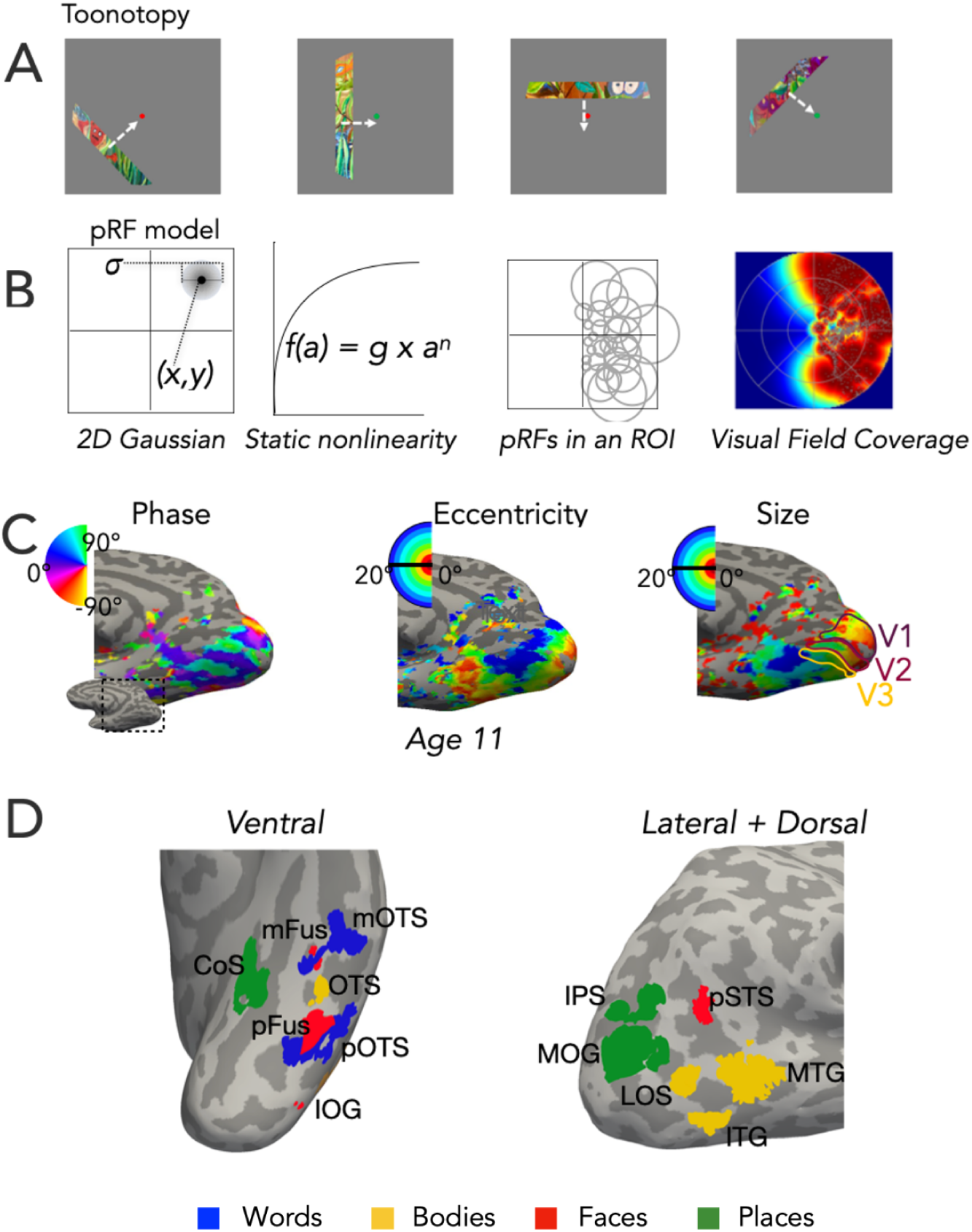
Experimental design and modeling. (A) Toonotopy stimuli from Finzi et al. (2021) features a bar with colorful cartoon images of faces, words, bodies, places, and objects that change at 8Hz sweeps across a gray background at 4 angles (0°, 45°, 90°, 135°) each in 2 directions. Participants fixated at the center dot and indicated when the dot changed colors. (B) Population receptive field compressive spatial summation (pRF CSS) model^17^. Left two panels show a single pRF with parameters of location (x,y) and size (*σ*) modeled by a 2D Gaussian followed by a compressive nonlinearity, used to model the response. Middle right panel shows schematic of the pRF distribution within an ROI, and the rightmost panel depicts the visual field coverage of all pRFs in left V1 ROI in an example 11 year old. (C) Phase, eccentricity, and size maps in an example adolescent (age 11) with V1 (purple), V2 (magenta), and V3 (gold) borders illustrated on the size map. All participants - Supplementary Figure 2 (D) Category selective ROIs in an example inflated left hemisphere of an 11 year old. ROIs are colored by category preference: *Green:* place-selective; *Red:* face-selective; *Blue:* word-selective; *Yellow:* body part selective. Acronyms: *CoS:* collateral sulcus; *mFus:* mid fusiform; *pFus:* posterior fusiform; *IOG:* inferior occipital sulcus; *OTS:* occipital temporal sulcus; *mOTS:* mid occipital temporal sulcus; *pOTS:* posterior occipital temporal sulcus; *IPS:* intraparietal sulcus; *MOG:* middle occipital gyrus; *LOS:* lateral occipital sulcus; *MTG:* middle temporal gyrus; *ITG:* inferior temporal gyrus; *pSTS:* posterior superior temporal sulcus; *IPS:* intraparietal sulcus; *MOG:* middle occipital gyrus.

From the Toonotopy experiment, we modeled each voxel’s pRF with a 2D Gaussian defined by four parameters – location in the visual field (*x,y*), size (*σ)*, and compressive nonlinearity (*n)*^17^ (**Fig. 1B**).

The Toonotopy experiment yielded the expected polar angle, eccentricity, and size maps with no qualitative differences across adolescents and adults (Fig. 1C). We found the standard polar angle maps with mirror reversals of the upper and lower visual field representations beginning in the calcarine sulcus (Fig. 1C, **Supplementary** Figure 2 **-all phase maps)**. Both adolescents and adults also exhibit the standard occipital eccentricity map, with a foveal to peripheral gradient extending anteriorly from the occipital pole, and a temporal eccentricity map, with a foveal to peripheral gradient extending from lateral to medial VTC (Fig. 1C**; Supplementary** Figure 3 **-all eccentricity maps**). Similarly, we observe the characteristic size maps with pRFs increasing from small to large moving anteriorly along the calcarine and from occipital cortex to VTC (Fig. 1C**; Supplementary** Figure 4 **-all size maps**). We observed no differences in pRF x-position, y-position, eccentricity, size, (**Supplementary Table 1**) or the relationship between size vs eccentricity in early visual cortex (V1, V2, V3) from adolescence to adulthood (**Supplementary Table 2; Supplementary** Figure 5).

From the category experiment, we defined category-selective regions of interest (ROIs) across all three streams (Fig. 1D). Face-selective ROIs were localized in the ventral (IOG-faces, pFus-faces, mFus-faces) and lateral (pSTS-faces) streams, body-selective ROIs in the ventral (OTS-bodies) and lateral (LOS-bodies, ITG-bodies, MTG-bodies) streams, place-selective ROIs in the ventral (CoS-places) and dorsal (MOG-places, IPS-places) streams, and word-selective ROIs in the ventral stream (pOTS-words, mOTS-words) (see Methods; **Supplementary Table 3**). Retinotopic maps extended to these ROIs in all participants (**Supplementary** Figure 2**-4**). As Toonotopy drives many of the voxels across high-level category-selective regions in both adolescents and adults (**Supplementary** Figure 1B**; Supplementary Table 4**), we then examined pRF properties in each ROI and participant.

### Across streams, pRF properties are differentially mature by adolescence

As the ventral stream contains selective regions for all four categories of interest and continues to mature during adolescence^9,35–38^, we first examined if the distributions of pRF properties in the ventral stream vary across adolescents and adults (*Left:* Fig. 2; *Right:* **Supplementary** Figure 6). In adolescents (Fig. 2A - top) and adults (Fig. 2A - bottom), pRFs centers are contralateral with left hemisphere pRFs in the right visual field and right hemisphere pRFs in the left visual field. Quantitatively, pRF horizontal (x) locations remain largely stable across development with distributions differing only in left pFus-faces (Fig. 2B, p = 7.3e-03, bootstrapped K-S-test with 10000 permutations), left CoS-places (Fig. 2B p = 7.1e-03, bootstrapped K-S-test), and right IPS-places (**Supplementary** Figure 7A, p= 0.0135, bootstrapped K-S-test) where pRFs are more contralateral in adolescents than adults (full stats in **Supplementary Table 5**). Distributions of pRF vertical (y) position also remain largely similar across adolescents and adults; pRFs are located significantly higher in the visual field in adolescents compared to adults only in left mFus-faces (Fig. 2C, p = 5.00e-05, bootstrapped K-S-test), right mFus-faces (**Supplementary** Figure 6B, p=0.044, bootstrapped K-S-test), and right mOTS-words (**Supplementary** Figure 6B, p = 5.30e-03, bootstrapped K-S-test). Finally, distributions of pRF sizes are similar across adolescents and adults for the majority of ventral category-selective regions. We observed significant differences across adolescents and adults in pRF sizes only left pOTS-words (Fig. 2D; p = .0003, bootstrapped K-S-test), left CoS-places (Fig. 2D; p = .0025, bootstrapped K-S-test), and right mFus-faces (**Supplementary** Figure 6C; p = .0008, bootstrapped K-S-test) whereby pRFs are larger in adolescents than adults.

**Figure 2.**
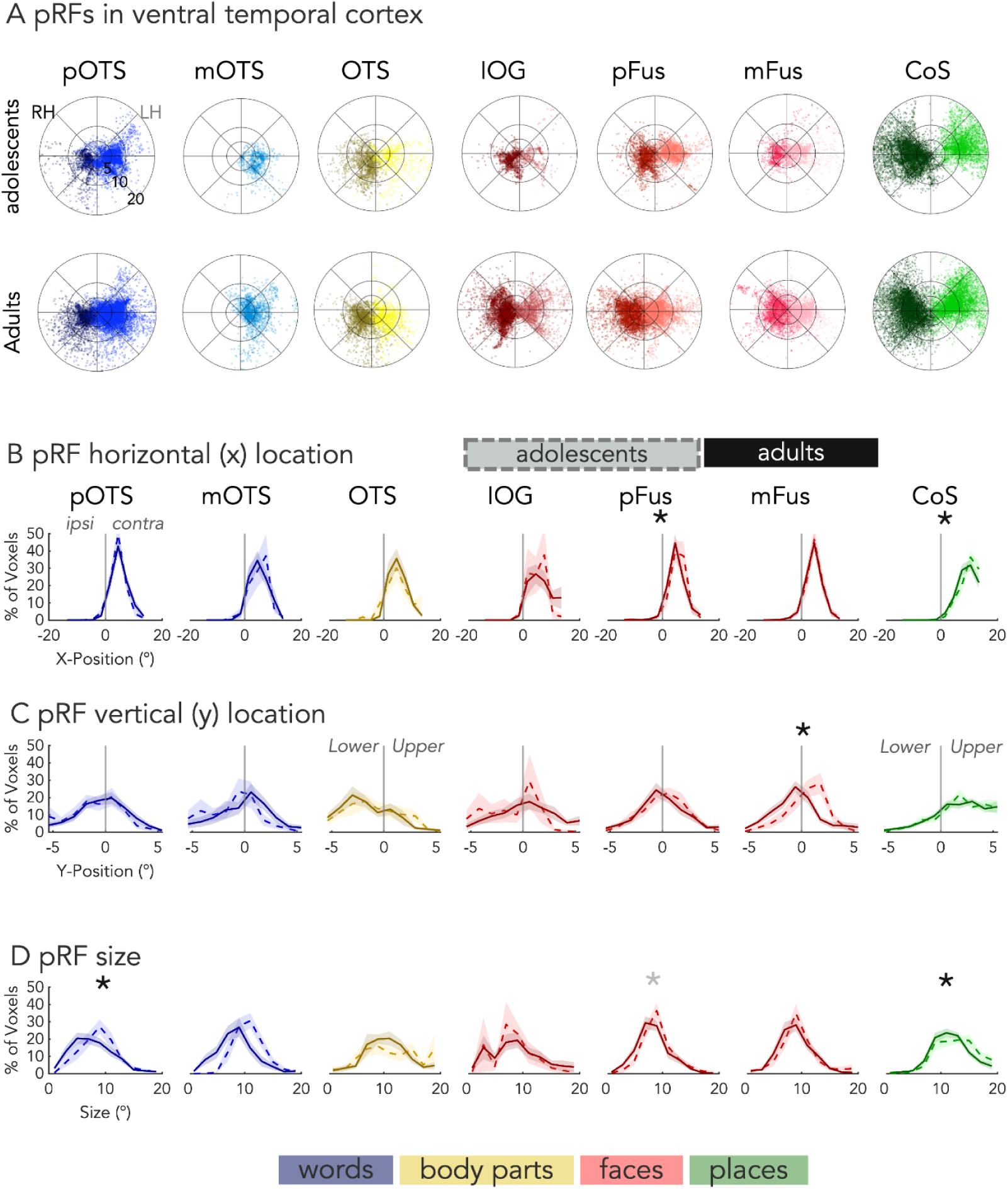
pRF properties in high-level category selective regions in the left ventral stream. (Right hemisphere – Supplementary Figure 6) (A) *Top:* pRF center polar plots for right hemisphere (dark) and left hemisphere (light) word-selective (blues; pOTS, mOTS), bodypart-selective (yellows; OTS), face-selective (reds; IOG, pFus, mFus), and place-selective (greens; CoS) regions in the ventral stream in adolescents ages 10 - 17. *Bottom:* pRF center polar plots same as A but in adults ages 22 - 32. (B) Distributions of x-position for left hemisphere ventral pRFs across the ipsilateral and contralateral visual fields for adults (solid) and adolescents (dashed). Black asterisk: significant difference between adolescents and adults using bootstrapped KS-tests with Bonferroni correction. Gray asterisk: significant difference between adolescents and adults using bootstrapped KS-tests before Bonferroni correction. (C) Same as (B) but for pRF y-position along the lower and upper visual fields. (D) Same as B but for pRF size. See Supplementary Table 3 for N’s per ROI.

To test if this pattern of development extends to other category-selective regions, we examined pRF properties in lateral and dorsal stream body, face, and place ROIs.. Like in the ventral stream, pRFs in lateral and dorsal ROIs are located in the contralateral visual field (*Left:* **Supplementary** Figure 7A**,B**, *Right:* **Supplementary** Figure 8A) with significant differences in distributions of pRF horizontal position only in right IPS-places (p = 0.0135, bootstrapped K-S-test with 10000 permutations) and no differences in vertical positions (*Left:* **Supplementary** Figure 7C, *Right:* **Supplementary** Figure 8B) across adolescents and adults (**Supplementary Table 5**). In the lateral stream, pRF sizes are large (∼10°– 20°; *Left*: **Supplementary** Figure 7D, *Right*: **Supplementary** Figure 8C) and are not significantly different across adolescents and adults (**Supplementary Table 5**). In the dorsal stream, we observed significant differences in the distribution of pRF sizes between age groups in bi-lateral IPS-places (Left: **Supplementary** Figure 7D, p = .0027, bootstrapped K-S-test; Right: **Supplementary** Figure 8C, p = .0002, bootstrapped K-S-test), which are larger in adolescents than adults.

Together, these data indicate that while pRFs properties are largely similar across adolescents and adults, there are some age-related differences, whereby a handful of ROIs exhibit pRFs that are larger and more peripheral in adolescents than adults.

### pRF vertical location varies more by category whereas pRF eccentricity and size vary more by stream

We next examined how pRF properties vary across categories and streams. Prior work suggests that pRF vertical (y) position and distance from the center of gaze (eccentricity) vary across categories^33^ and streams^34^, but these predictions have yet to be tested across multiple streams and categories. To test these hypotheses, we compared mean pRF vertical position, eccentricity, and size across body-, face-, and place-selective ROIs in all three streams. Using linear mixed-effects models (LMMs; see Methods), we tested if mean pRF properties vary across stream, category, hemisphere, and age group.

pRF vertical position significantly varies across ROIs with different category preferences (Fig. 3A, main effect of category: F(1,693.18) = 54; p = 1.69e-22, LMM - Methods, eq. 1) and does not significantly vary across adolescents and adults (full stats - **Supplementary Table 6**). Indeed, pRFs in body-selective ROIs (OTS, LOS, ITG, MTG) have a lower visual field bias, pRFs in face-selective regions (IOG, pFus, mFus, pSTS) show no vertical bias, and pRFs in left place-selective CoS and IPS have an upper visual field bias (Fig. 3A). Additionally, we observed hemispheric (category x hemi: F(1, 683.74 = 3.347, p = 0.033; LMM) and stream differences (category x stream: F(2, 684.25) = 12.146, p = 6.56e-6) in vertical bias. In body-selective ROIs, the lower field bias is larger in the right than left hemisphere. In place-selective ROIs the vertical bias varies across streams and hemispheres, with pRFs in bilateral MOG showing a lower visual field bias and pRFs in right CoS- and IPS-places showing no vertical bias.

**Figure 3.**
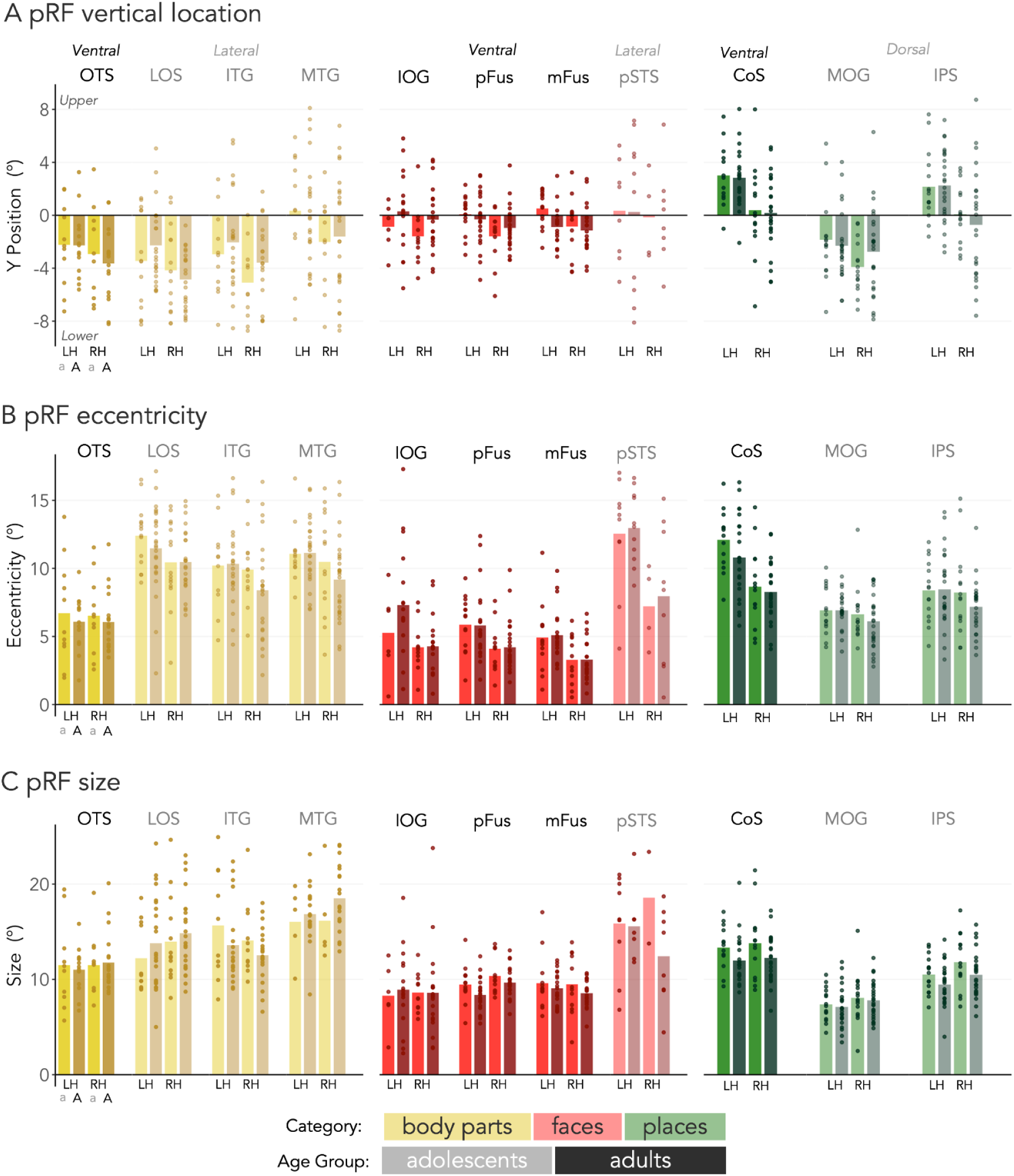
pRF vertical position, eccentricity, and size across categories, streams, and age groups. (A) Bars indicate average pRF y-position for each age group.Bar plots for body-selective ROIs (yellow), face-selective ROIs (red), and place-selective ROIs (green). For each category, lighter bars are adolescents and darker colors are adults. Saturation indicates stream: *more saturated:* ventral; *less saturated:* lateral/dorsal. *Dots:* individual participants. *LH:* left hemisphere; *RH:* right hemisphere. (B) Same as A but for pRF eccentricity. (C) Same as A but for pRF size. See Supplementary Table 3 for N’s per ROI.

By contrast, pRF eccentricity varies significantly by stream (Fig. 3B, F(1,693.18) = 84.616; p = 4e-9, LMM - Methods, eq. 1) and stream by category (F(2,684.25) = 106.505; p = 5.379e-41, LMM), with no significant differences across age groups (full stats, **Supplementary Table 7**). Across adolescents and adults, pRFs in ventral body- and face-selective regions are more foveal than pRFs in lateral body- and face-selective ROIs, which extend to the periphery (Fig. 3B). Conversely, pRFs in ventral place-selective ROIs are more peripheral than pRFs in dorsal place-selective ROIs (Fig. 3B). We also observe an interaction of stream, category, and hemisphere whereby pRFs in left pSTS-faces and left CoS-places are more peripheral than in their right hemisphere counterparts (stream x category x hemisphere: F(2, 682.72 = 10.701; p = 2.654e-5, LMM).

Like pRF eccentricity, pRF sizes vary significantly by stream (Fig. 3C, F(1,694.48) = 35.637; p = 3.79e-9, LMM) and stream by category (F(2,684.82) = 25.67; p = 1.79e-11, LMM; full stats in **Supplementary Table 8**). pRFs in ventral body- and face-selective regions are smaller than in their lateral counterparts, whereas pRFs in ventral place-selective ROIs are larger than their dorsal counterparts (Fig. 3C). Different from other pRF properties, pRF size varies significantly across adolescents and adults (main effect of age: F(1,64.2) = 5.623; p = 0.021, LMM) with adolescents having larger pRFs than adults. This age group difference is most pronounced in right pSTS-faces and in bilateral place-selective ROIs (age x stream x category x hemi: F(2,683.16) = 3.615; p = 0.027).

Overall, these data indicate that mean pRF vertical position varies predominantly by category whereas pRF eccentricity and size vary primarily by stream and additionally by category. Furthermore, pRF properties differentially develop, as pRF vertical position and eccentricity are largely mature by adolescence yet pRF size continues to develop from adolescence into adulthood.

### Visual field coverage continues to develop during adolescence

To examine how pRF properties of individual voxels impact the total region of the visual field processed by each visual region, we calculated the visual field coverage (VFC) of pRFs spanning each ROI (*Left:* Fig. 4A; *Right:* **Supplementary** Figure 9A). We quantified each ROI’s VFC as its total full-width at half-maximum (FWHM; Fig. 4A - blue dashed circle) – measured from the center of mass across all its pRFs (CoM; Fig. 4A - blue asterisk).

**Figure 4.**
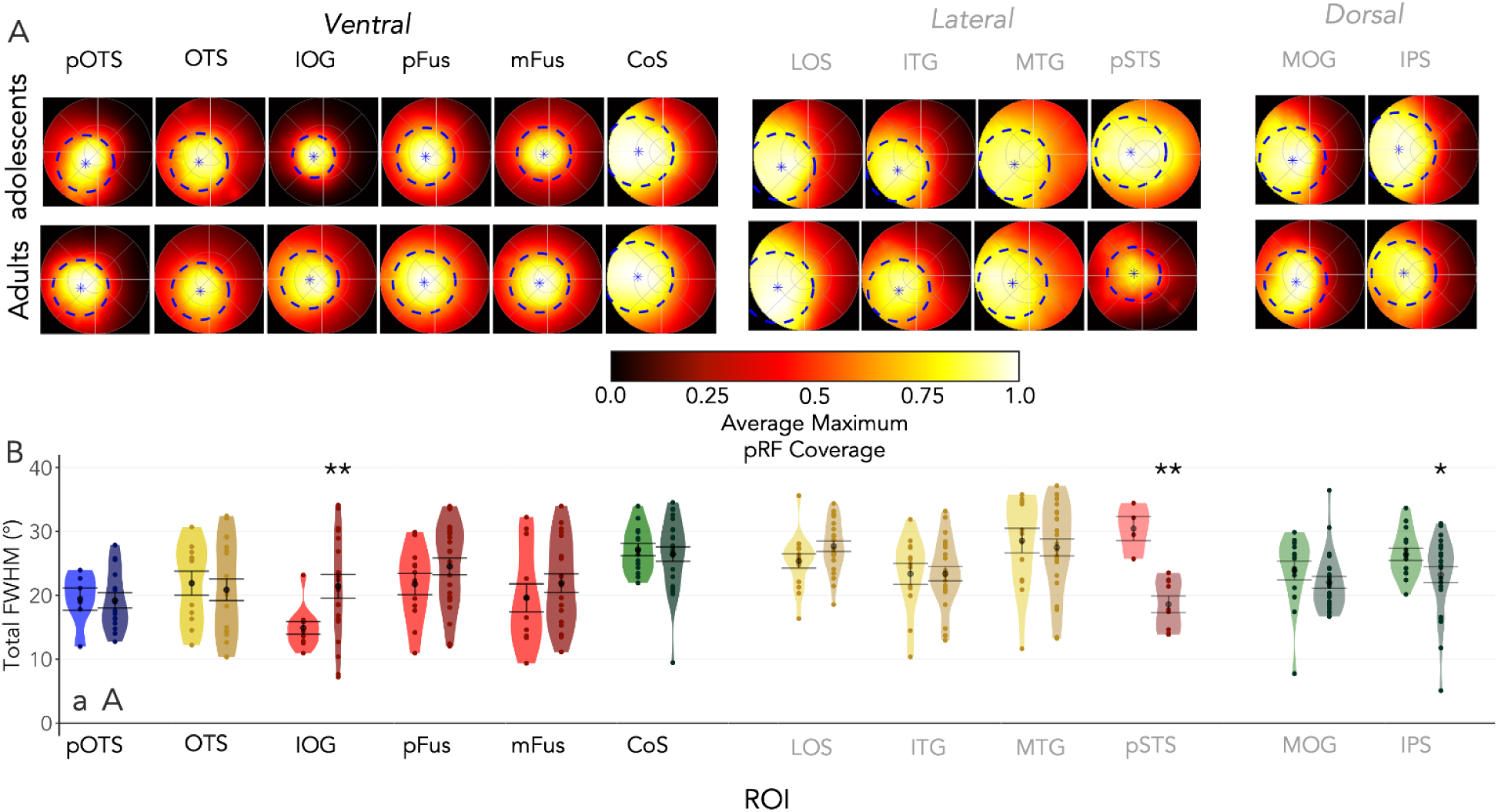
Visual field coverage (VFC) of right hemisphere category-selective ROIs in adolescents and adults (Left hemisphere – Supplementary Figure 9). (A) Average VFC of each ROI in the right hemisphere across adolescents (top row) and adults (bottom row) hemisphere with a warm gradient ranging from higher coverage in white and lower coverage in black. VFC is calculated as the maximum response of pRFs covering each point in the visual field for each participant and is then averaged across participants in the group. *Blue asterisks:* average center of mass of the VFC. *Blue dotted lines:* average full-width half max (FWHM) of the coverage. (B) Violin plots of total FWHM in visual degrees for category-selective ROIs (colors as in Figure 3) in the right hemisphere ventral, lateral, and dorsal streams in adolescents (a; lighter colors) and adults (A; darker colors). *Black circle:* mean FWHM. *Error bars:* ± SE. See Supplementary Table 3 for N’s per ROI.

VFC retains the category- and stream-specific differences observed for individual pRF properties. With respect to category, the coverage is largely the central visual field in face- and word-selective ROIs, the lower visual field in body-selective ROIs, and the upper visual field in place-selective ROIs (*Left:* Fig. 4A**,B**; *Right:* **Supplementary** Figure 9A**,B**). With respect to stream, ventral body and face ROIs exhibit more central coverage (∼30% of FWHM in the center 5° of the visual field), while dorsal-lateral ROIs show more peripheral coverage (just ∼15% of the FWHM in the central 5°). However, coverage extends into the periphery in both ventral and dorsal place-selective ROIs (∼13% of FWHM in center 5°). To quantify these observations, we used a LMM to test if VFC varies by age group, stream, category, and hemisphere and another ventral LMM to test if the VFC varies across age group, category, and hemisphere within the ventral stream (Methods, eq. 1,2). Indeed, VFC varies significantly by stream and category (stream: F(1, 694.72) = 7.024, p = 0.008, category: F(2,686.50) = 4.094, p = 0.017, *LMM*; F(3, 358.41) = 13.369, p = 1.914e-08, *ventral LMM;* full stats **Supplementary Table 9)**.

VFC also exhibits significant differential development from adolescence into adulthood (age group x stream x category: p = 2.62*10^-3^, F(2, 685.28) = 6; *LMM*; age group x stream x hemisphere: p = 7.68*10^-3^, F(1, 693.29) = 7.15; *LMM*; **Supplementary Table 9**). This development is driven by significant decreases in the VFC in right pSTS-faces (post-hoc t-test: t(27.06) = 5.20, p = 1.98*10^-3^) and right IPS-places (post-hoc t-test: t(37.99) = 2.06, p = 0.05), as well as significant increases in coverage in right IOG-faces (post-hoc t-test: t(6.04) = −3.09, p = 4.61*10^-3^) from adolescence to adulthood (*Left:* Fig. 4B; *Right:* **Supplementary** Figure 9B). However, the ventral stream shows no significant development in VFC (*ventral LMM*; **Supplementary Table 9**).

### Category-selectivity develops during adolescence, independent from visual field coverage

As prior work finds that ventral category-selective ROIs change in selectivity and consequently size from adolescence to adulthood^9,35,43^, we next examined if the size of category-selective ROIs across all streams continues to develop during adolescence. Thus, we quantified the size of category-selective ROIs and tested if ROI size varies by stream, category, hemisphere, and age group using LMMs (Methods, eq. 3-4; **Supplementary Table 10**).

We observe differential development of category-selective ROI size (Fig. 5; category x group: F(2,777.97) = 5.906, p = 0.0028; *LMM;* F(3,385.84) = 4.286, p = 0.0054; *ventral LMM*). Several face- and word-selective ROIs as well as lateral body-selective ROIs increase in size, but several place-selective ROIs decrease in size from adolescence into adulthood (Fig. 5 **- asterisks**, betas - **Supplementary Table 11**). In a complementary analysis, we also examined mean selectivity (t-value) in independent 10mm disk ROIs centered on each category-selective ROI and found that mean selectivity also develops during adolescence (**Supplementary Table 12**).

**Figure 5.**
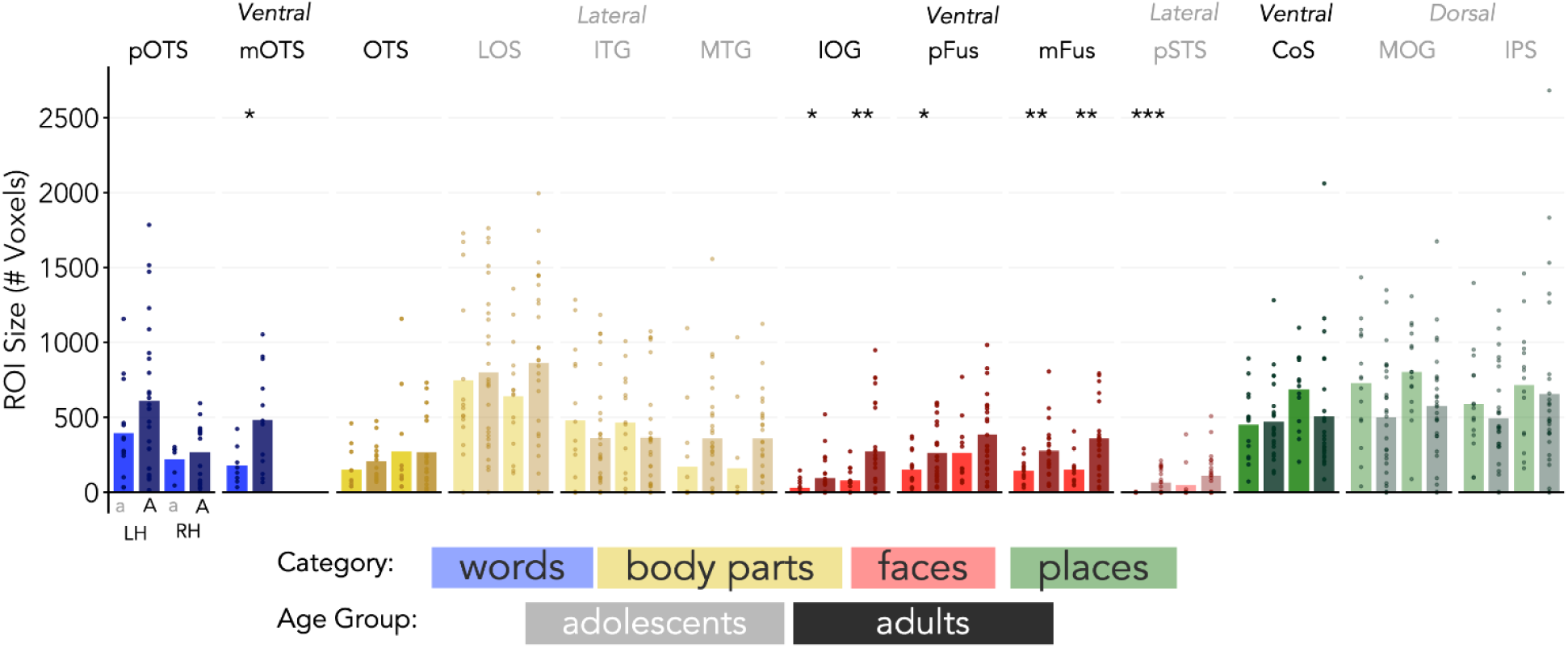
Category-selective ROI size in adolescents and adults. Bar plots of average category-selective ROI sizes. For word-selective ROIs (blue), body-selective ROIs (yellow), face-selective ROIs (red), and place-selective ROIs (green). For each category, lighter bars are adolescents and darker colors are adults. Saturation indicates stream: *more saturated:* ventral; *less saturated:* lateral/dorsal. *Dots:* represent individual participants. *LH:* left hemisphere; *RH:* right hemisphere.See Supplementary Table 3 for N’s per ROI.

We used LMMs to test if development in the size (and selectivity) of category selective ROIs is linked to the development of pRFs and VFC (**Supplementary Figure 10A, B;** Methods, eq. 5-6). We observe a consistent link between category selectivity and VFC: larger ROI size is linked to larger VFC (F(1,241.05) = 53.509, p = 6.53*10^-12^; β = 21.813; **Supplementary Table 13, S14**) and greater category selectivity is also linked to larger VFC (F(1,233.13) = 24.865, p = 1.201*10^-6^; β = 0.077; **Supplementary Table 13, S14**). Larger ROI size is also significantly associated with greater eccentricity (F(1,239.46) = 4.442, p =0.036; β = 12.263; **Supplementary Table 13, S14**). Interestingly, these relationships do not differ by age group: age does not explain additional variance in ROI size or mean t-value (F’s < 0.493, p’s > 0.487; **Supplementary Table 13, S14**). Together, these results show that category selectivity is coupled with spatial integration by pRFs, and this relationship is not mediated by age group.

## Discussion

Here, we examined how pRF properties and category selectivity develop from adolescence to adulthood in category-selective regions spanning the ventral, dorsal, and lateral visual streams Two central insights emerge. First, high-level visual cortex continues to develop from adolescence into adulthood. Second, its functional organization is multidimensional: pRF properties and selectivity jointly vary by category, stream, and hemisphere, rather than being governed by a single organizing principle.

### Functional fingerprints of category-selective regions across streams

To summarize the data, we characterize per age group each ROI’s functional fingerprint across multiple dimensions – pRF y-position, eccentricity, size, FWHM, normalized ROI size (Fig. 6).

**Figure 6.**
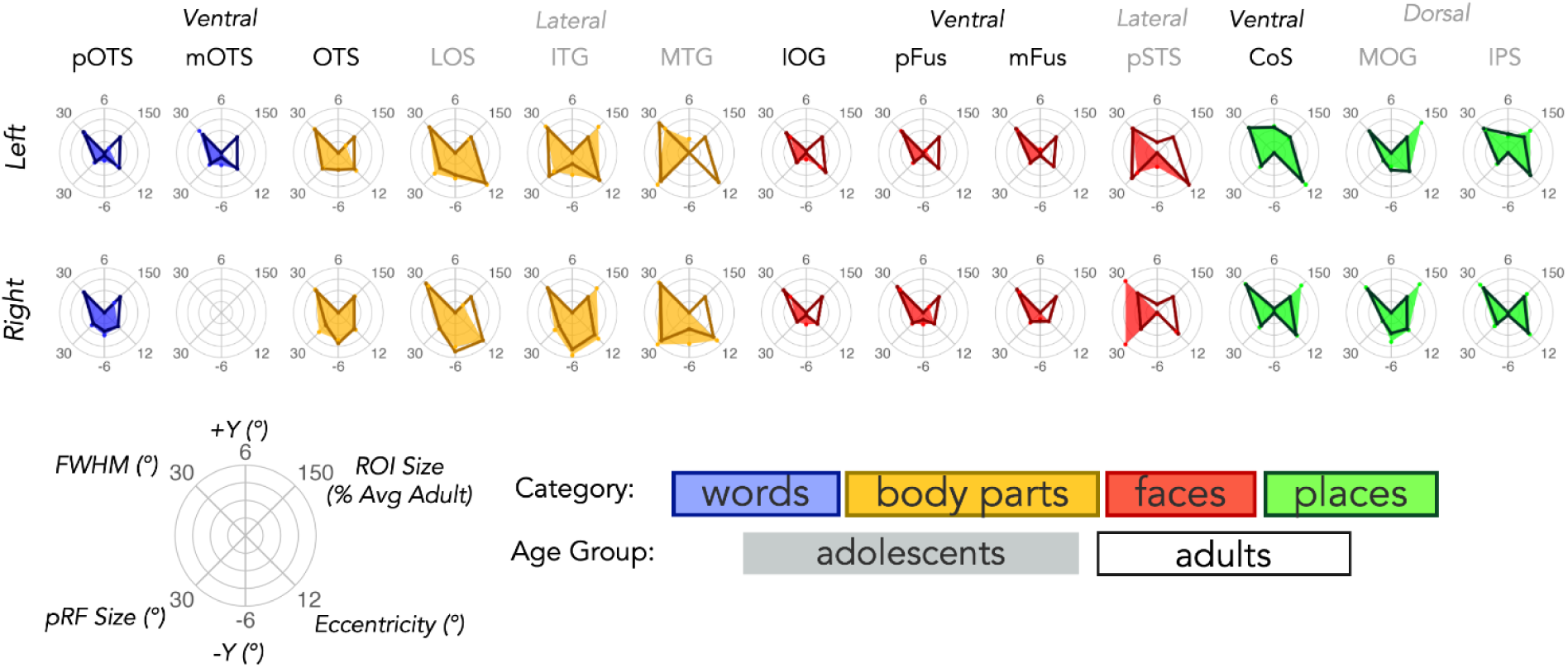
Functional fingerprint of category-selective ROIs. Top: Left hemisphere fingerprint plots of average positive y-position (90°), ROI size (45°; normalized to the average adult size for that ROI; see Methods), eccentricity (−45°), negative y-position (−90°), pRF size (−135°), and total FWHM (135°) for word - (blue), body - (yellow), face - (red), and place-selective (green) ROIs in the ventral, lateral, and dorsal streams. *Adolescents:* shaded; *Adults:* outline. *Top:* Left hemisphere; *Bottom:* Right hemisphere. See Supplementary Table 3 for N’s per ROI.

Functional fingerprints reveal systematic differences across categories, streams, and hemispheres, underscoring that no single feature captures the functional characteristics of high-level visual regions or their developmental trajectories during adolescence.

The pattern of results summarized by the functional fingerprint challenges accounts that emphasize a single dominant functional organizing feature of high-level visual cortex, such as eccentricity biases linked to category preference^26,27,30^ or vertical position biases linked to streams^33,34^. Instead, our data indicate that pRF eccentricity, size, and visual field coverage vary across both category preference and stream, while pRF vertical position varies primarily with category preference and hemisphere. Specifically, face- and body-selective regions show smaller eccentricity, smaller pRF size, and more central visual field coverage in the ventral than the lateral stream. In contrast, place-selective regions show larger eccentricity, larger pRFs, and broader coverage in the ventral than the dorsal stream. Additionally, vertical pRF biases differ by category and hemisphere: body-selective regions show a lower-field bias especially in the right hemisphere, place-selective regions show an upper-field bias selectively only in the left hemisphere, and face- and word-selective regions do not have a vertical bias. Together, these findings emphasize that the functional fingerprints of regions in high-level visual cortex are jointly determined by category preference, stream, and hemisphere.

### A mechanistic hypothesis: category and task specific visual spatial processing

We hypothesize that the functional fingerprint of high-level visual cortex reflects the convergence of category-specific visual experience and stream-specific task demands. Specifically, the statistical regularities with which observers view particular categories while performing specific visual tasks may shape pRF properties and consequently visuospatial processing across development to become optimized for processing behaviorally relevant information in particular regions of the visual field^20,22,25–28,30,44–47^.

For example, face recognition and reading, tasks that are associated with the ventral stream^22,35,48,49^, require high spatial acuity associated with foveal vision. Consistent with this demand, pRFs (and visual field coverage) in ventral face- and word-selective regions are centered around fixation and have relatively small spatial integration windows^26,30,50,51^. In contrast, tasks associated with the lateral stream, like processing dynamic information necessary for action and social recognition, may require integrating information across larger areas of the visual field including the periphery. Correspondingly, pRFs in lateral face- and body-selective regions are larger and more peripheral than those in the ventral stream. Likewise, scene recognition, a task associated with the ventral stream, may require integration over larger portions of the visual field than navigational affordances^44,52–54^, a task associated with the dorsal stream, paralleling the larger pRFs and wider visual field coverage in ventral than dorsal place-selective regions. Additionally, regularities in viewing stimuli from different categories may also generate pRF vertical biases. For example, frequent fixation on faces will more often put bodies in the lower visual field^28^. These observations support a mechanistic account in which category- and task-specific visual demands shape pRF properties in high-level visual cortex, emphasizing the role of visual experience in the development of functional profiles of these category-selective regions.

### A timeline of functional development in high-level visual cortex

Prior work suggests that pRF properties in early visual cortex are largely mature by ages 5–7, whereas pRFs in ventral temporal and lateral occipitotemporal cortex continue to develop during childhood^25,39–41^. Extending this developmental timeline, we find that by adolescence, pRF location – horizontal and vertical position as well as eccentricity – is largely adult-like in most category-selective regions. In contrast, pRF size and visual field coverage show more protracted development from adolescence to adulthood. Several ROIs exhibit decreases in pRF size and/or visual field coverage (e.g., left pFus-faces, right mFus-faces, right pSTS-faces, left pOTS-words, and bilateral IPS-places), whereas others show increased coverage (e.g., right IOG-faces). These changes indicate that while pRF location is largely mature by adolescence, the window of spatial integration either at the level of a voxel or at the level of the population code across a visual region^11,21,22,25,28,55,56^ continues to develop during adolescence.

Category selectivity also differentially develops during adolescence. Face- and word-selective regions increase in selectivity and expand in size whereas place-selective regions and lateral body ROIs show decreased selectivity and shrink in size. Thus, we find that category selectivity has a prolonged development during adolescence which involves both increases and decreases in selectivity. These data are consistent with longitudinal data showing recycling of limb-selectivity to face- and word-selectivity in the ventral stream from childhood to adolescence within individual children^9,20,35^, suggesting that cortical recycling^51,56^ may continue into adulthood and is not just limited to the ventral stream. Together, these results emphasize adolescence as a period of continued functional plasticity, extending the window for potential intervention for developmental disorders associated with high level visual cortex like autism^57,58^, dyslexia^38,59,60^, and developmental prosopagnosia^61–64^.

### Implications for how experience during development may shape high-level visual cortex

Our category-by-task hypothesis is consistent with two non-exclusive developmental trajectories. One possibility is that pRF properties are largely established early in development, and category selectivity emerges later as a result of statistical regularities in viewing particular categories during specific tasks^20,22,25–28,30,44–47,65^. Another possibility is that streams have coarse pRF properties early in development that are progressively refined by category and task specific experience, such that visual experience simultaneously shapes both category selectivity and pRF properties to support task-relevant spatial computations across the visual field.

Although the present data do not provide information about the relationship between visual experience and either the development of pRFs or category selectivity, they identify developmental windows when pRFs in high-level visual cortex are malleable (pre-adolescence) and when spatial integration and category selectivity are malleable (pre-adulthood). Future longitudinal studies in children can further elucidate functional development by examining rich functional fingerprints across multiple dimensions beyond category-selectivity and pRF properties such as texture^55^, animacy^66^, object size^67^, format^68–71^, affordances^44,72^, social information^6^, and dynamics^73^ together with measurements of children’s visual experience using mobile eye tracking during naturalistic tasks. These types of data will be crucial for testing how children’s visual experience and task demands shape functional and pRF properties across regions and streams.

## Conclusions

Overall, our findings delineate the developmental trajectory of high-level visual cortex that extends from adolescence into adulthood and reveal that pRF organization across ventral, lateral, and dorsal streams is inherently multidimensional. Functional fingerprints reveal that each category-selective region follows its own developmental path, driven not by a single feature but by the joint maturation of pRFs, spatial integration, and category selectivity. This multidimensional perspective provides a principled framework for understanding how high-level visual cortex develops and becomes functionally organized.

## Materials and Methods

### Participants

19 neurologically typical adolescents aged 10 to 17 years old (M = 13.74 ± 2.13; 11 females, 8 males) and 27 adults (ages 22 - 32; M = 25.52 ± 3.00; 13 females, 14 males) participated in this study. 4 of the adolescents participated in two scanning sessions approximately a year apart. All participants had normal or corrected-to-normal vision. Participants, or their parents, gave written informed consent, and all procedures were approved by the Stanford Internal Review Board on Human Subjects Research.

Sessions were excluded from the analysis if within scan motion and/or between scan motion was greater than 2 voxels. Of the 19 adolescent sessions, 4 participants were excluded based on these motion criteria. No adult sessions were excluded. Of the remaining participants, there were no significant differences in motion between adolescents and adults. After exclusion, we analyzed the data of 15 adolescents (ages 10 - 17; 9 females, 6 males) and 27 adults (ages 22 - 32; 13 females, 14 males).

### Data Acquisition

#### MRI

Participants were scanned using a General Electric Discovery MR750 3T scanner located in the Center for Cognitive and Neurobiological Imaging (CNI) at Stanford University. A phase-array 32-channel head coil was used for the category localizer experiment and to obtain anatomical scans. For the toonotopy experiment, a 16 channel head coil was used.

#### Anatomical scans

For each participant, we obtained a whole-brian anatomical volume using a T1-weighted pulse sequence (TI = 450ms, 1×1×1mm, flip angle = 12 degrees, FoV = 204 mm). Anatomical images of each brain were used for segmentation of the gray/white matter boundary.

#### Toonotopy

Participants completed four runs of a wide-field pRF mapping fMRI experiment with cartoon stimuli, which we refer to as Toonotopy^20^. In the experiment (Fig. 1A), bars of width 5.7° swept a circular 40°x40° (visual angle) aperture with a fixation dot at center. The bars swept the visual field at four orientations (0°, 45°, 90°, 135°) in eight directions (2 opposite directions orthogonal to each orientation). The cartoon stimuli randomly changed at a rate of 8 Hz with blanks (mean luminance gray background with fixation) appearing at regular intervals. During each run, participants fixated on the central dot and were instructed to press a button whenever the dot changed colors. Each run was 3 minutes and 24 seconds long. Average performance on the color changing fixation task was 97% in adolescents and 93.14% in adults.

#### Category experiment

The same participants underwent an fMRI category experiment, which is used to identify voxels whose neural response is stronger to one category vs. many other categories^42^. In each run, participants were presented with stimuli from five domains, each with two categories (Fig. 2A; faces: child, adult; bodies: whole, limbs; places: corridors, houses; objects: cars, guitars; characters: words, numbers). Images within the same category were presented in 4s blocks at a rate of 2Hz and were not repeated across blocks or runs. 4s blank trials were also presented throughout a block. During a run, each category was presented eight times in counterbalanced order, with the order differing for each run. Throughout the experiment, participants fixated on a central dot and performed an oddball detection task, pressing a button when phase-scrambled images randomly appeared. Each participant completed 3 runs of the category localizer experiments with different images; each run was 5 minutes and 18 seconds long.

### Data Analysis

#### Anatomical data analysis

T1-weighted images were automatically segmented using FreeSurfer (FreeSurfer 7.0.0:^74^) and then manually validated and corrected using ITKGray. Cortical reconstructions were generated from these segmentations using FreeSurfer.

#### fMRI data analysis

Data were processed and analyzed in MATLAB using mrVista (http://github.com/vistalab). All data were analyzed within the individual participant native brain space. Functional data were manually aligned to the T1-weighted volume. The manual alignment was then optimized using robust multiresolution alignment. For participants with more than one session, functional data were aligned to the anatomical scan taken closest to the date of the functional scan. Data were not spatially smoothed and were restricted to the cortical ribbon. Functional data were motion-corrected within and between scans using mrVista motion correction algorithms. Quality assurance was also performed to determine exclusions based on motion.

#### mrVista to FreeSurfer conversion

To visualize functional maps and draw regions of interest (ROIs), eccentricity and phase maps from Toonotopy and category selectivity maps from the category localizer experiment were converted from mrVista to FreeSurfer coordinates and projected onto the inflated cortical surface reconstruction for each individual participant in Freeview (FreeSurfer 7.2.0).

#### Defining ROIs

ROIs were drawn using Freeview (FreeSurfer 7.2.0) on the inflated cortical surface reconstruction of each participant’s brain then projected back to mrVista for analysis. ROIs were defined by JKY and JOC.

##### V1-V3

Using polar angle and eccentricity maps from the Toonotopy experiment thresholded at 20% variance explained, we defined early retinotopic visual areas (V1, V2, V3; Fig. 1B) in both hemispheres in each individual. Boundaries between retinotopic areas were defined as the middle of polar angle reversals at the horizontal or vertical meridian representations, and each area included foveal to peripheral representations^13,18^. Dorsal (V1d, V2d, V3d) and ventral (V1v, V2v, V3v) components of each visual area were defined separately and combined in analysis to create representations of the entire visual field (V1, V2, V3).

##### Category-selective regions of interest (ROIs)

From the category experiment, statistical contrast maps of each category domain versus all other category domains (i.e. faces > all other stimuli) were thresholded at a *t-value* > 3 at the voxel level, as in previous experiments^20,25,42^. Using these contrast maps, category-selective ROIs in the ventral, lateral, and dorsal streams were defined in each subject as clusters of voxels selective for a category located at a particular anatomical landmark (Fig. 2B).

Face-selective voxels (contrast: adult and child faces > all other categories) were defined in the inferior occipital gyrus (IOG-faces), posterior fusiform gyrus (pFus-faces), mid fusiform gyrus (mFus-faces), and posterior superior temporal sulcus (pSTS-faces). Word-selective voxels (contrast: words > all other categories except numbers) were defined in the posterior occipital temporal sulcus (pOTS-words) and mid occipital temporal sulcus (mOTS-words). Because we could identify only in a minority of subjects (∼20%) the right mOTS-words, mOTS-words was only defined in the left hemisphere. Bodypart-selective voxels (contrast: bodies and limbs > all other categories) were defined in the occipital temporal sulcus (OTS-bodies), lateral occipital sulcus (LOS-bodies), inferior occipital gyrus (IOG-bodies), and mid temporal gyrus (MTG-bodies). Place-selective voxels (contrast: corridors and houses > all other categories) were defined in the collateral sulcus (CoS-places), intraparietal sulcus (IPS-places), and mid occipital gyrus (MOG-places^75,76^).

Because we were only able to identify pSTS-face and MTG-bodies in a minority of adolescents (pSTS-faces: left 0/15; right 3/15; MTG-bodies: left: 5/15, right 4/15) we used group probability maps of these ROIs from adults projected to individual cortical surfaces for both adolescents and adults. The probability maps were thresholded at voxels found in 30% or more of the participants.

After defining the category ROIs in each individual, only ROIs that contained ten or more voxels with at least 20% variance explained during the Toonotopy experiment were included in subsequent analyses (see **Supplementary Table 2** for number of subjects per ROI). Ventral and dorsal stream ROIs had a higher proportion of voxels with >20% variance explained during the Toonotopy experiment (∼80%) compared to lateral stream ROIs (∼20-60%) consistently across age groups and hemispheres (**Supplementary** Figure 1B).

Overall, we report data from 7 category-selective ROIs in the ventral stream (IOG-faces, pFus-faces, mFus-faces, pOTS-words, mOTS-words, OTS-bodies, CoS-places), 4 ROIs in the lateral stream (pSTS-faces, LOS-bodies, ITG-bodies, MTG-bodies), and 2 ROIs in the dorsal stream (MOG-places, IPS-places). As prior research combined the dorsal and lateral stream into a single dorsal stream^23,30,34^ and because streams differ in the number and type of category-selective regions – e.g., dorsal stream contains only one category (places) – throughout this study we compare the ventral stream and a combined dorsal-lateral stream.

#### Estimating pRFs

The time-course data of the Toonotopy experiment were transformed from functional slices to the T1-weighted whole brain anatomy using trilinear interpolation. The pRF of each voxel was modeled using the compressive spatial summation (CSS) model^17^ (Fig. 2B) using VISTA lab software (http://github.com/vistalab). A pRF is modeled independently for every voxel by fitting a 2D Gaussian with a center (x,y) and a size determined by the standard deviation (σ) of the Gaussian in degrees of visual angle. An exponent parameter (0≤n≤1) is additionally fit for each voxel to capture the compressive nonlinearity of pRFs. pRF size is thus defined as α/√n. x, y, and σ are iteratively optimized to minimize the root mean squared error between the observed and predicted time-series. Eccentricity (√x^2^+y^2^) and phase (αtan(y/x)) of each voxel were derived from the center (x,y) coordinates and used to generate eccentricity and phase maps, respectively. As in prior studies^20,22^ we report pRF parameters for voxels in which the variance explained by the pRF model is greater than 20%.

##### pRF size vs. eccentricity

To evaluate the relationship between pRF size and eccentricity (Fig. 1), all analyzed voxels in each participant’s ROI were entered into a linear regression comparing pRF size to eccentricity. A line of best fit was derived for each participant, and the slope and intercept of the line was averaged across participants of each group (Fig. 1D).

#### Visual field coverage (VFC)

VFC, the region of the visual field processed by the set of pRFs spanning an ROI, was calculated for each ROI and participant. Each voxel’s pRFs were represented by a 2D Gaussian centered on their centers (x,y) with size σ, and voxels were included if their variance explained exceeded 20%. To create individual VFC maps, we used the maximum profile method and computed at each point in the visual field the strongest pRF response from any voxel in the ROI. To create group VFC maps (Fig. 4A, B), we averaged maps across participants within each group (adolescents, adults). Each average VFC map thus reflects the average maximal spatial extent of the ROI’s visual field representation.

##### Estimating the full-width half-max (FWHM) of the VFC

For each ROI and participant, we calculated the FWHM (Fig. 4A, B - black dashed line), which provides a standardized measure of the spatial extent of the VFC by estimating the diameter, in visual degrees, of the cross section of the VFC in which it reaches half of its maximum amplitude. The FWHM was determined by fitting a circular Gaussian centered at the center of mass, namely, the peak response (Fig. 4A, B - white asterisk), of the VFC, using the equation: *FWHM = 2√2*ln(2)√σ* where σ is the standard deviation of the Gaussian.

#### Category selectivity analysis

To examine the development of category selectivity in our functional category ROIs, we quantified the mean t-value and the ROI size for each ROI in each individual and then compared across age groups.

##### Mean t-value

To evaluate how the strength of category selectivity develops, we quantified the mean t-value, which indicates how strongly an ROI responds to one visual category versus all other categories. To control for ROI size differences across participants, groups, and ROIs and focus on selectivity, we generated a 10mm radius disk ROI centered on the original functional category ROI. The 10mm disk ROI approximated the average ROI size across all participants and regions. We then calculated the mean t-value (unthresholded) of the 10mm disk ROI for the category and ROI was drawn for every participant.

##### ROI size

To examine the extent of cortex involved in processing each category and how this develops from adolescence into adulthood, we measured ROI size defined as the the number of category-selective voxels with t-value > 3. This measurement was done for each category ROI in every participant. To assess differences between adults and adolescents in Figure 6, ROI size was normalized within each ROI and hemisphere such that the adult mean ROI size was set to 100%, and adolescent values were expressed relative to this baseline.

### Statistical Analysis

All statistical analyses were conducted using R version 4.2.2. Except for analysis of EVC, which included all subjects, the number of participants included in each statistical test, based on whether or not they had an ROI, was consistent across all analyses and can be found in **Supplementary Table 3**.

#### KS-Tests

To compare differences in pRF distributions between adolescents and adults within an ROI, we conducted two-sample Kolmogorov-Smirnov (KS) tests with bootstrapped resampling for each individual ROI and pRF parameter (x, y, size). We performed 10,000 bootstrap iterations, in which we drew an equal number of voxels from all adolescents and all adult for each ROI. The number of voxels drawn was determined by taking the average number of voxels included in an ROI across both adolescents and adults. In each iteration, we recomputed the KS-test using a stream-specific Bonferroni threshold (Ventral: p < 0.0038; Lateral/Dorsal: p < 0.0042). We report the adjusted mean p-value, the mean KS-statistic across all bootstraps and the 95% CI. The p-value reported from the bootstrap is 1 minus the fraction of significant rejections across iterations (**Supplementary Table 5**).

#### Linear Mixed Modeling (LMM)

LMMs were conducted using the lme4^77^ and emmeans packages in R (https://CRAN.R-project.org/package=lme4, https://CRAN.R-project.org/package=emmeans). Dependent variables included pRF parameters (x-position, y-position, size, eccentricity), visual field coverage (FWHM), and category selectivity (mean t-value, ROI size). LMMs were used to accommodate fixed and random effects, repeated measures, and incomplete data (e.g., not all participants had all ROIs). Primary fixed effects included age group (adolescents, adults), stream (ventral, dorsal-lateral), category (faces, words, bodies, places), and hemisphere (left, right), with participant as a random effect to control for inter-subject variability.

We use LMMs to quantify significant differences of factors of interest. In the first analysis, referred to as *LMM*, we compared metrics of interest across age groups, streams, categories, and hemispheres for face, body, and place ROIs, which are distributed across multiple streams. As prior research combines the dorsal and lateral stream into a single dorsal stream and because streams differ in the number and type of category-selective regions, e.g., dorsal stream contains only one category (places), we compare the ventral stream and a combined dorsal-lateral stream.

(1) *pRF property ∼ **stream * category** * group * hemisphere + (1 | participant)*

In the second analysis, referred to as *ventral LMM,* we assessed differences across age group, category, and hemisphere for ROIs within the ventral stream in order to include word-selective ROIs, which were only identified in the ventral stream.

(2) *pRF property ∼ **category** * group * hemisphere + (1 | participant)*

We used LMMs to test if ROI size and category selectivity (mean t-value) change significantly across age group, streams, categories, and hemispheres, with participant as a random effect.

(3) *ROI Size (selectivity) ∼ stream x category x group x hemisphere +(1 | participant)*

We also used LMM to test within the ventral stream if ROI size and category selectivity vary significantly across age group, category, and hemisphere, with participant as a random effect.

(4) *ROI Size (selectivity) ∼ category x group x hemisphere +(1|participant)*

In the last set of LMMs we tested if there is a significant relationship between ROI size and category selectivity and pRF properties, with participant as a random effect:

(5) *ROI Size ∼ VFC FWHM + pRF Size + pRF eccentricity + group + (1 | participant);*

(6) *Mean T-Value ∼ VFC FWHM + pRF Size + pRF eccentricity + age group + (1 | participant).*

Full results and statistics from all LMMs are in **Supplementary Tables 1-14**. Additional analyses with age as a continuous variable are also included in the **Supplementary Tables 6-10, 12.**

#### Post-hoc testing

Post-hoc two-tailed t-tests were performed using the t.test function in R to further examine significant main effects of age group and age group interactions from the LMMs. For each ROI, t-tests compared the mean value of the dependent variable between adolescents and adults. Post-hoc t-tests were conducted for pRF size, total FWHM, category selectivity (mean t-value), and ROI size, given main effects of or interactions with age group. Only significant post-hoc t-tests are reported in the main text.

## Data Availability

Raw data available upon request. Processed data available on Github: https://github.com/VPNL/toonCat

## Code Availability

fMRI data were analyzed using the open source mrVista software package (https://github.com/vistalab/vistasoft). Custom code for processing the pRF experiment and functional localizer, reproducing figures, and statistics can be found at https://github.com/VPNL/toonCat.

## Author Contributions

J.K.Y. analyzed the data and wrote the manuscript. J.C. contributed to data analysis and contributed to the manuscript. D.F. designed the experiment, collected data, and contributed to the manuscript. K.G-S. designed the experiment, contributed to data analysis, and wrote the manuscript.

## Acknowledgements

This material is based upon work supported by the National Science Foundation Graduate Research Fellowship Program under Grant No. (DGE-2146755) awarded to J.K.Y., NIH grants (grant numbers RO1EY022318, RO1EY023915 to K.G.S.), a William R. and Sara Hart Kimball Stanford Graduate Fellowship awarded to D.F., and a Symbolic Systems Summer Internship Program internship (J.C.).

