## Supplementary Figures and Tables for "From adolescence to adulthood: functional fingerprints of high-level visual cortex reveal differential development of visuospatial processing"

**Supplementary Information**

**Figures**

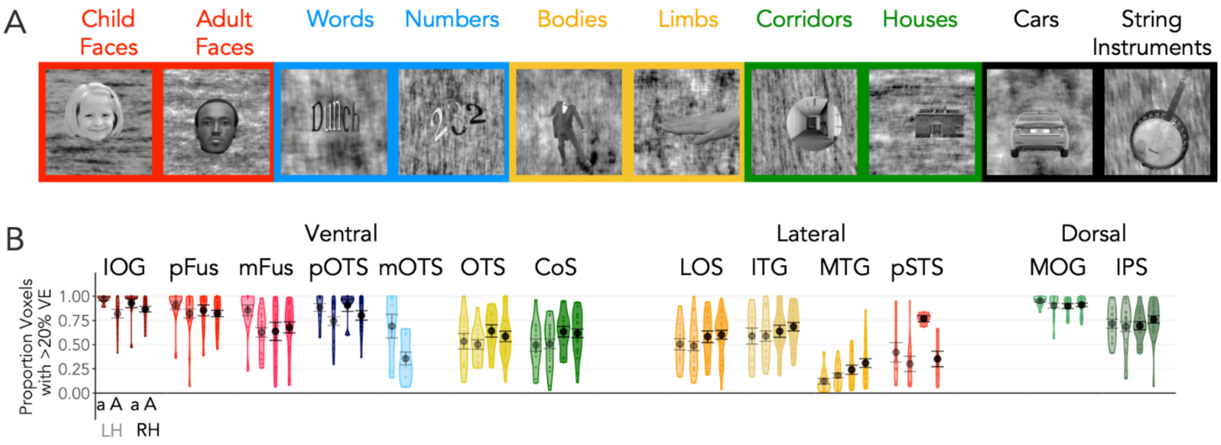

**Supplementary Figure 1. Category-selective regions are modulated by the Toonotopy experiment.**

(A) Category Experiment: Example stimuli of faces (children, adults; red), characters (words, numbers; blue), bodies and limbs (yellow), places (corridors, houses; green), and objects (car, guitar; black) from the category experiment. (B) Violin plots of proportion of voxels with greater than 20% variance explained by a pRF model in the Toonotopy experiment in category-selective ROIs in ventral, lateral, and dorsal streams in adolescents (a) and adults (A). *Light colors:* left hemisphere; *Darker colors:* right hemisphere. ) Error bars: ± SE (standard error of the mean). Each dot is a participant.

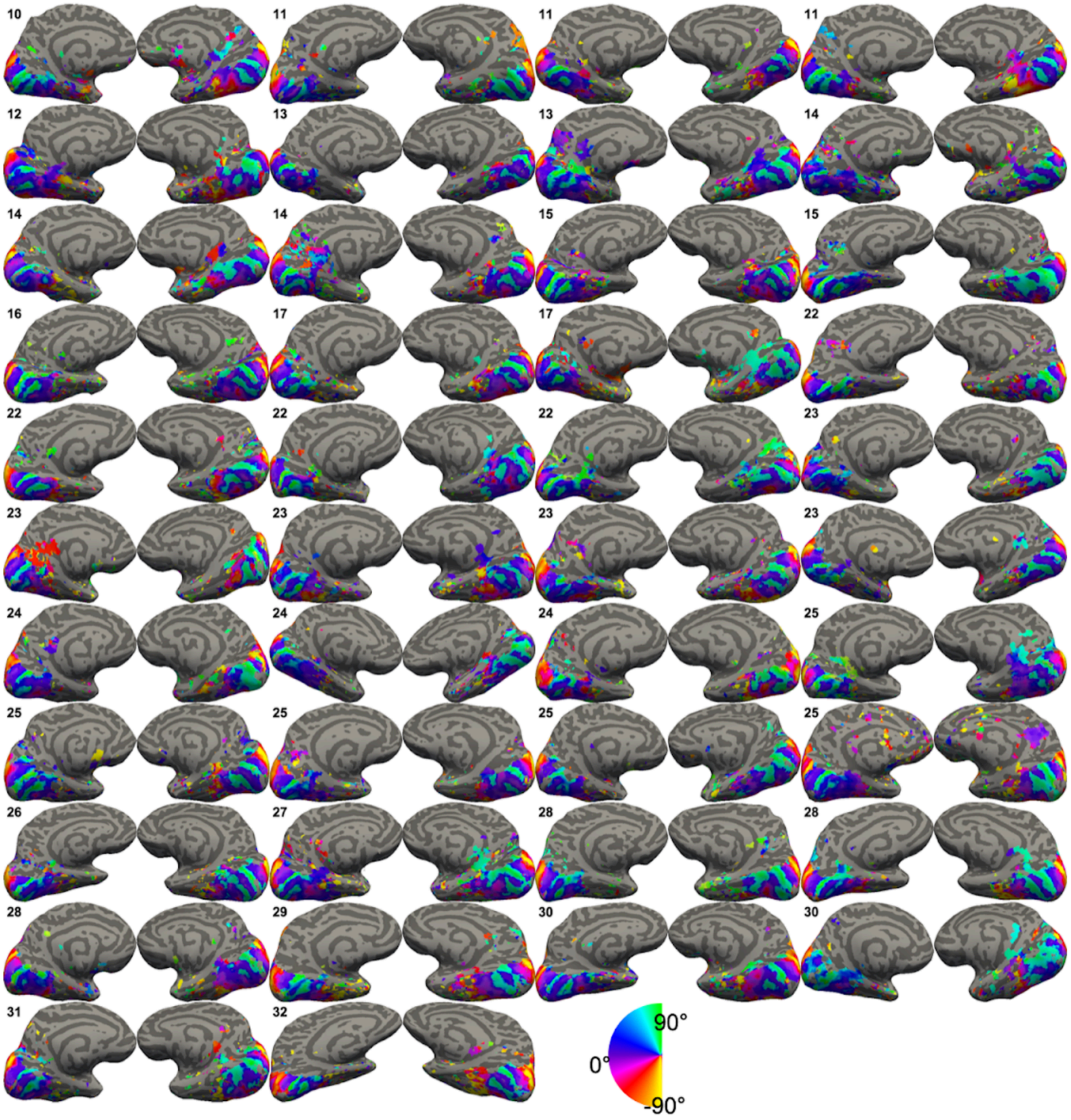

**Supplementary Figure 2. Phase maps for all participants.**

Phase maps for each individual participant thresholded at 20% variance explained in the Toonotopy experiment on left and right hemisphere medial inflated surfaces. All analyses were done within each participant's brain. Each pair of panels (left, right hemisphere) shows one participant, organized from youngest (left) to oldest (right) within each row. Age is indicated in black at the top left corner of each pair of hemispheres. Color wheel on bottom right indicates pRF phase in visual degrees.

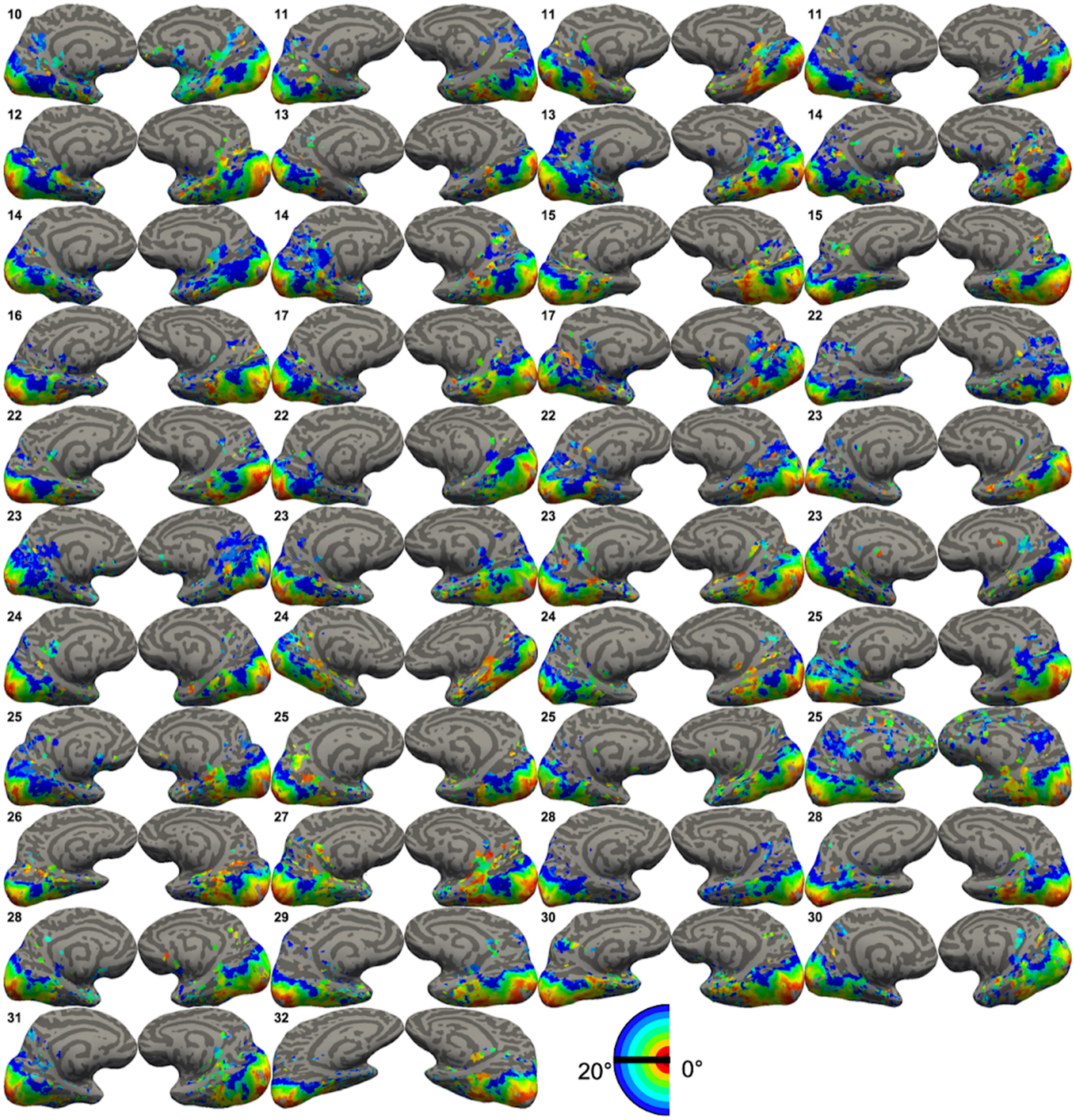

**Supplementary Figure 3. Eccentricity maps for all participants.**
Eccentricity maps for each individual participant thresholded at 20% variance explained in the Toonotopy experiment on left and right hemisphere medial inflated surfaces. All analyses were done within each participant's brain. Each pair of panels (left, right hemisphere) shows one participant, organized from youngest (left) to oldest (right) within each row. Age is indicated in black at the top left corner of each pair of hemispheres. Color wheel indicates pRF eccentricity in visual degrees.

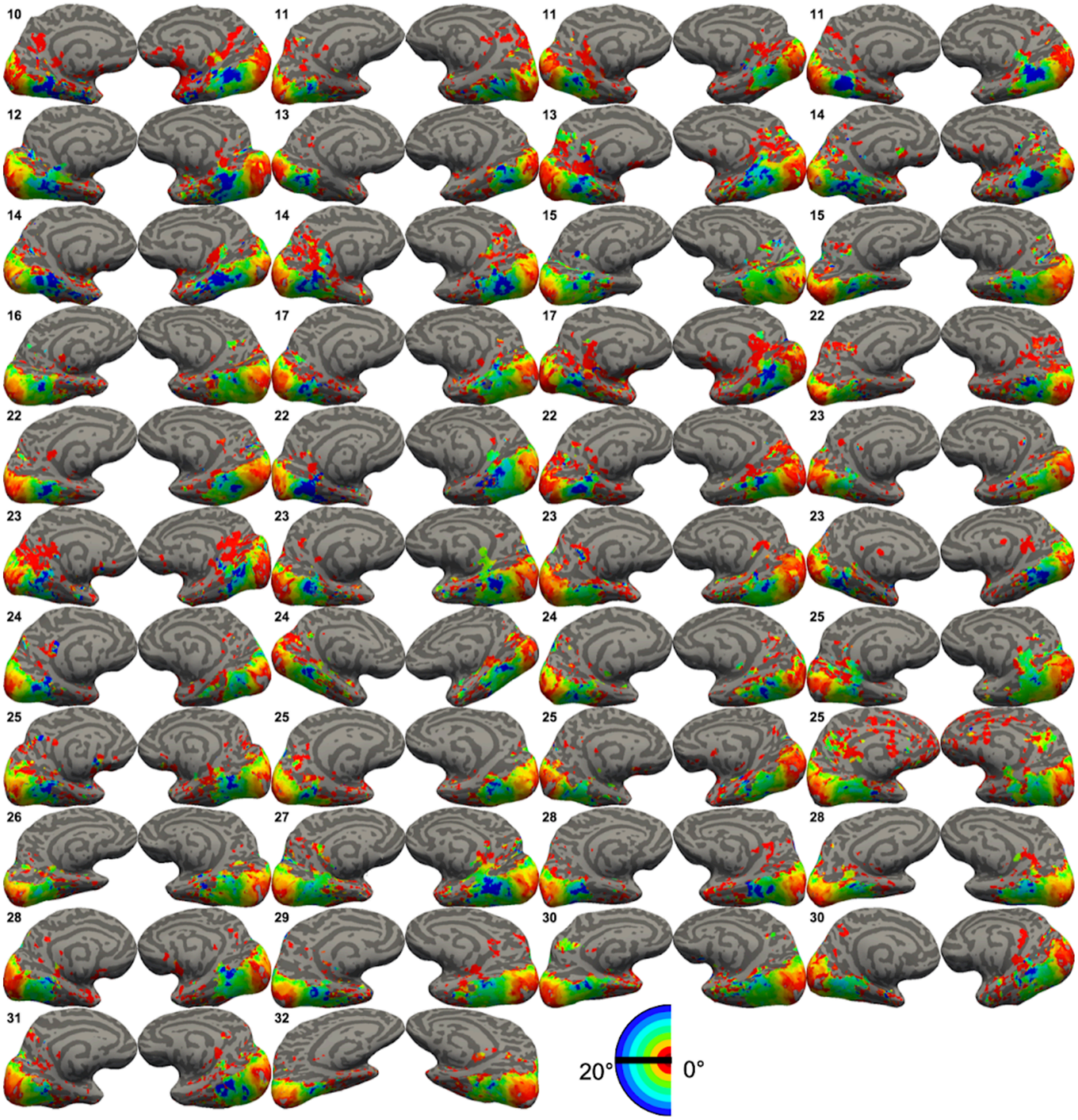

**Supplementary Figure 4. Size maps for all participants.**
Size maps for each individual participant thresholded at 20% variance explained in the Toonotopy experiment on left and right hemisphere medial inflated surfaces. All analyses were done within each participant's brain. Each pair of panels (left, right hemisphere) shows one participant, organized from youngest (left) to oldest (right) within each row. Age is indicated in black at the top left corner of each pair of hemispheres. Color wheel indicates pRF size in visual degrees.

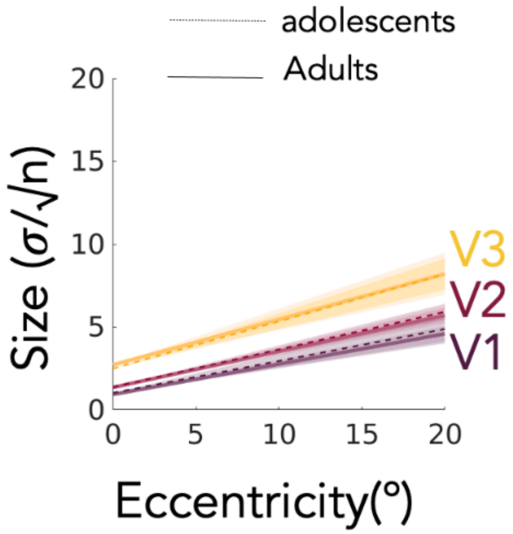

**Supplementary Figure 5. Relationship between pRF size vs eccentricity in early visual cortex**

Data show relationship between pRF size vs eccentricity across all adolescents and adults for V1, V2, and V3. Lines indicate average, and shaded areas indicate confidence interval. Data show that pRF size versus eccentricity relationship is similar across adolescents (dotted line) and adults (solid line) in early visual cortex (V1 - V3).

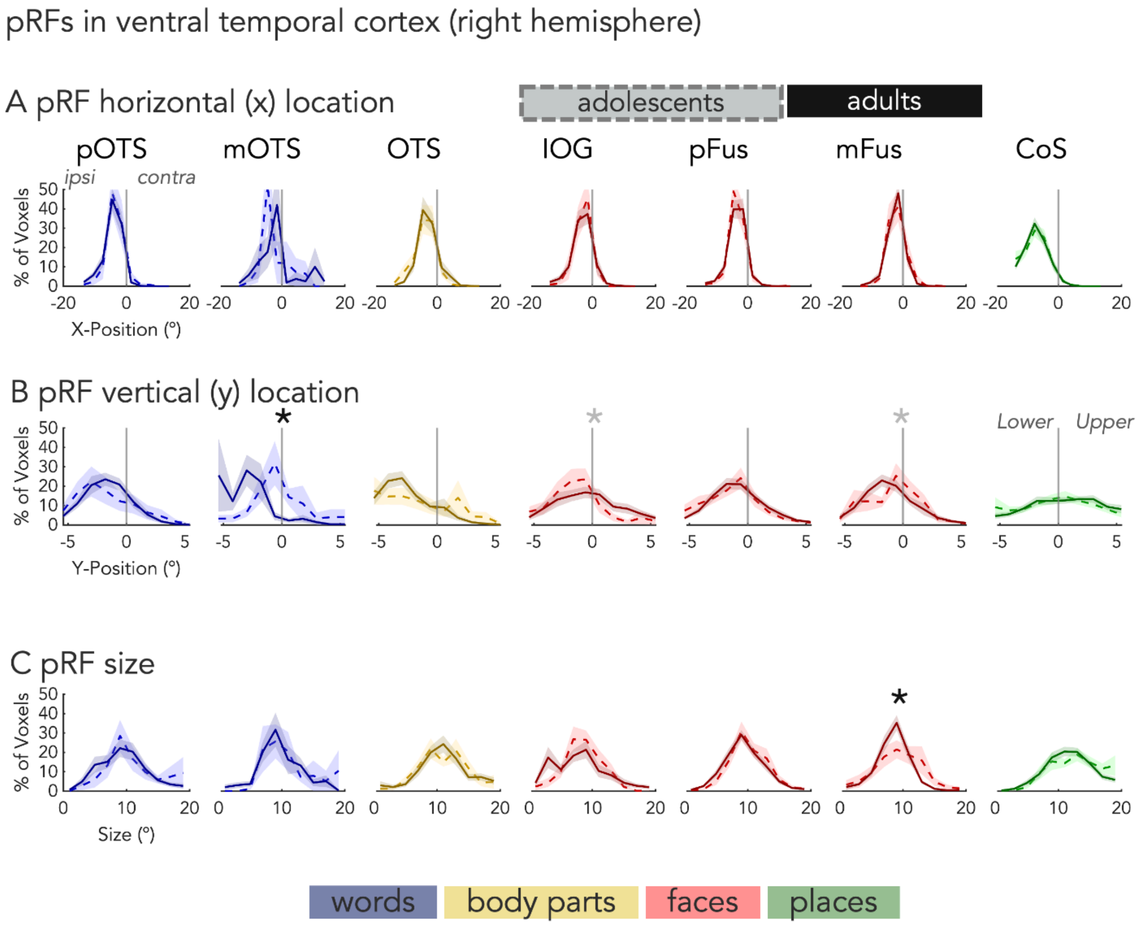

**Supplementary Figure 6. pRF properties in high-level category selective regions in the right ventral stream.**

(A) Distributions of x-position for right hemisphere ventral pRFs across the ipsilateral and contralateral visual fields for adults (solid) and adolescents (dashed). *Black asterisk*: significant difference between adolescents and adults using bootstrapped KS-tests with Bonferroni correction. *Gray asterisk:* significant difference between adolescents and adults using bootstrapped KS-tests before Bonferroni correction. (B) Same as (A) but for pRF y-position along the lower and upper visual fields. (C) Same as (A) but for pRF size.

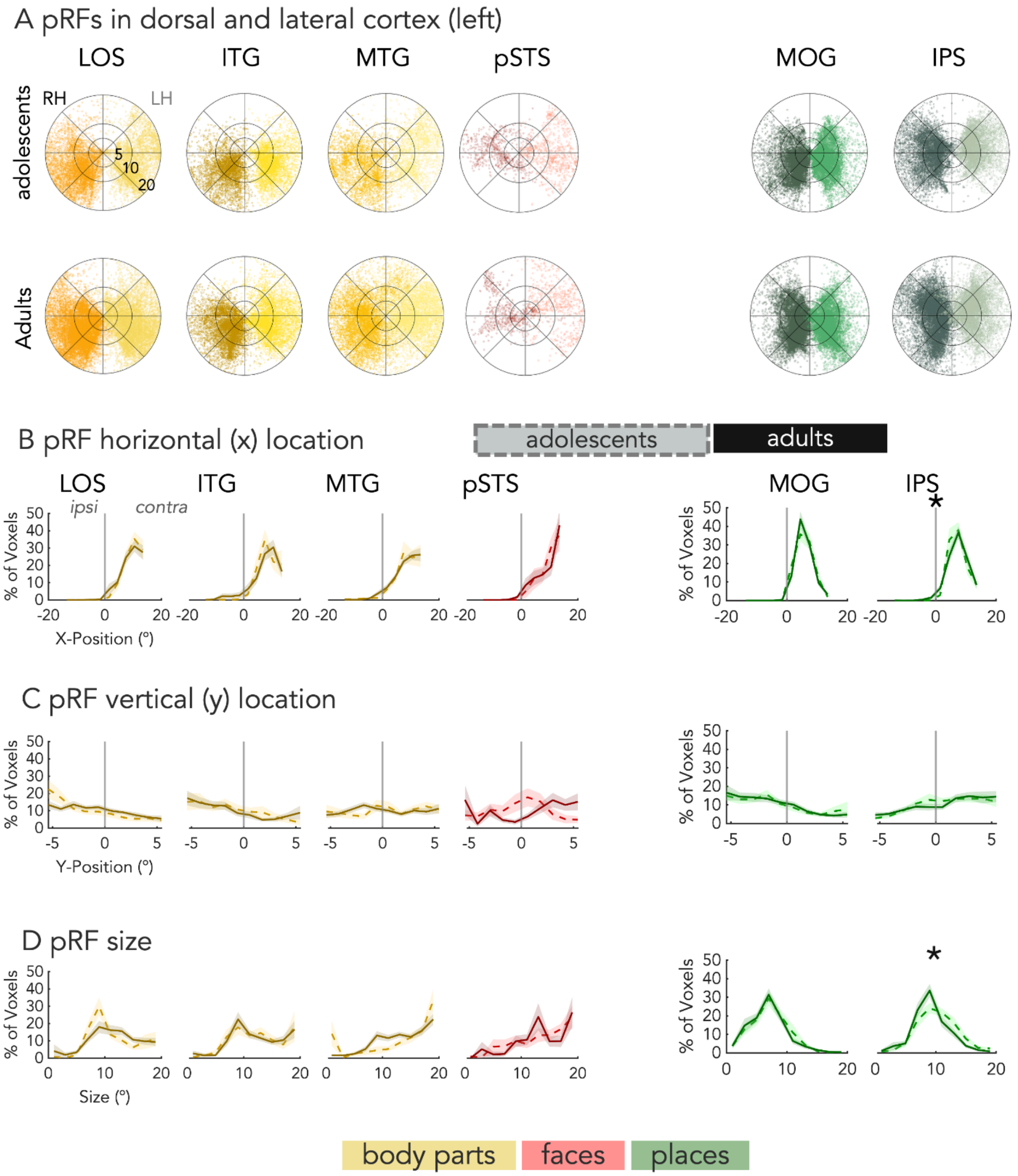

**Supplementary Figure 7. pRF properties in high-level category selective regions in the left lateral and dorsal streams.**

(A) *Top:* pRF center polar plots for right hemisphere (dark) and left hemisphere (light) bodypart-selective (yellows; LOS, ITG, MTG), face-selective (red; pSTS), and place-selective (greens; MOG, IPS) regions in the ventral stream in adolescents ages 10 - 17. *Bottom:* pRF center polar plots same as A but in adults ages 22 - 32. (B) Distributions of x-position for left hemisphere lateral and dorsal pRFs across the ipsilateral and contralateral visual fields for adults (solid) and adolescents (dashed). *Black asterisk:* significant difference between adolescents and adults using bootstrapped KS-tests with Bonferroni correction. *Gray asterisk:* significant difference between adolescents and adults using bootstrapped KS-tests before Bonferroni correction. (C) Same as (B) but for pRF y-position along the lower and upper visual fields. (D) Same as B but for pRF size.

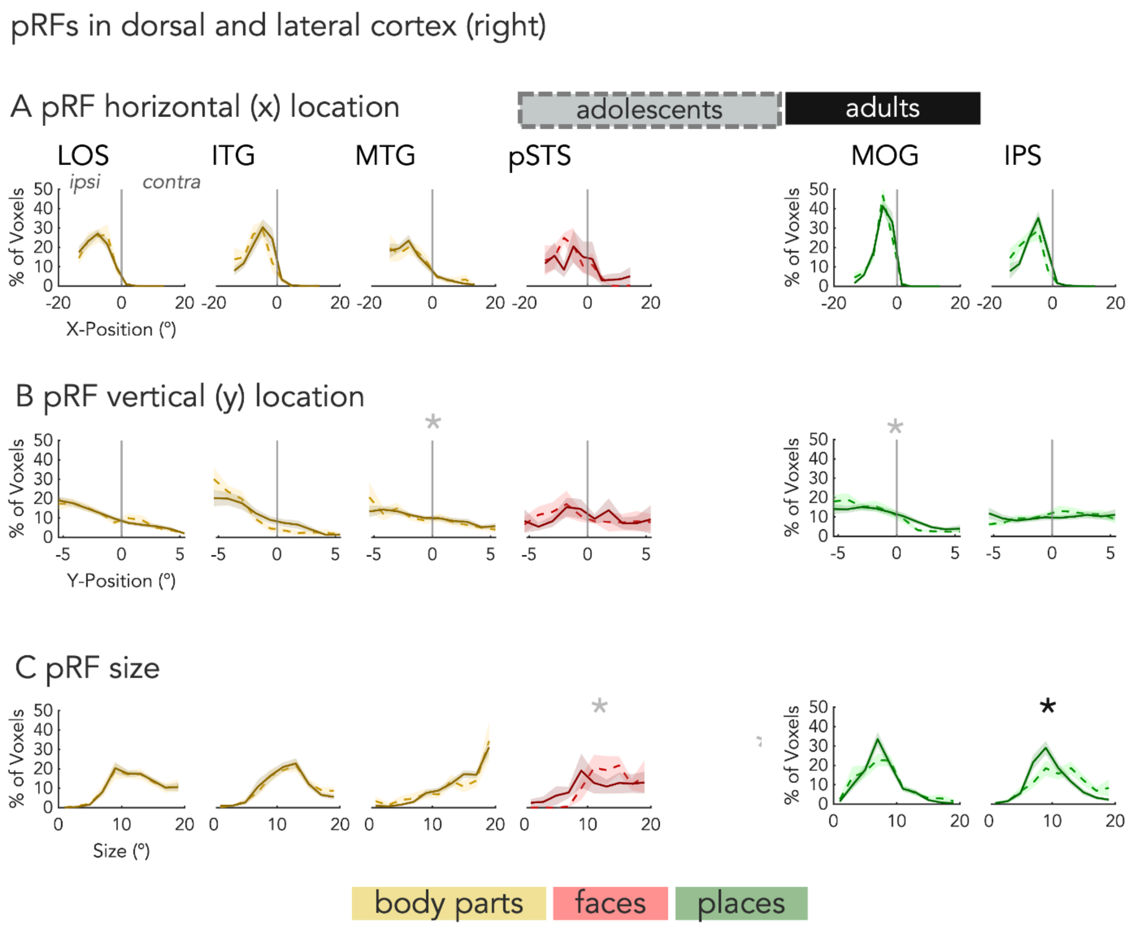

**Supplementary Figure 8. pRF properties in high-level category selective regions in the right lateral and dorsal streams.**

(A) Distributions of x-position for right hemisphere lateral and dorsal pRFs across the ipsilateral and contralateral visual fields for adults (solid) and adolescents (dashed). *Black asterisk:* significant difference between adolescents and adults using bootstrapped KS-tests with Bonferroni correction. *Gray asterisk:* significant difference between adolescents and adults using bootstrapped KS-tests before Bonferroni correction. (B) Same as (A) but for pRF y-position along the lower and upper visual fields. (C) Same as (A) but for pRF size.

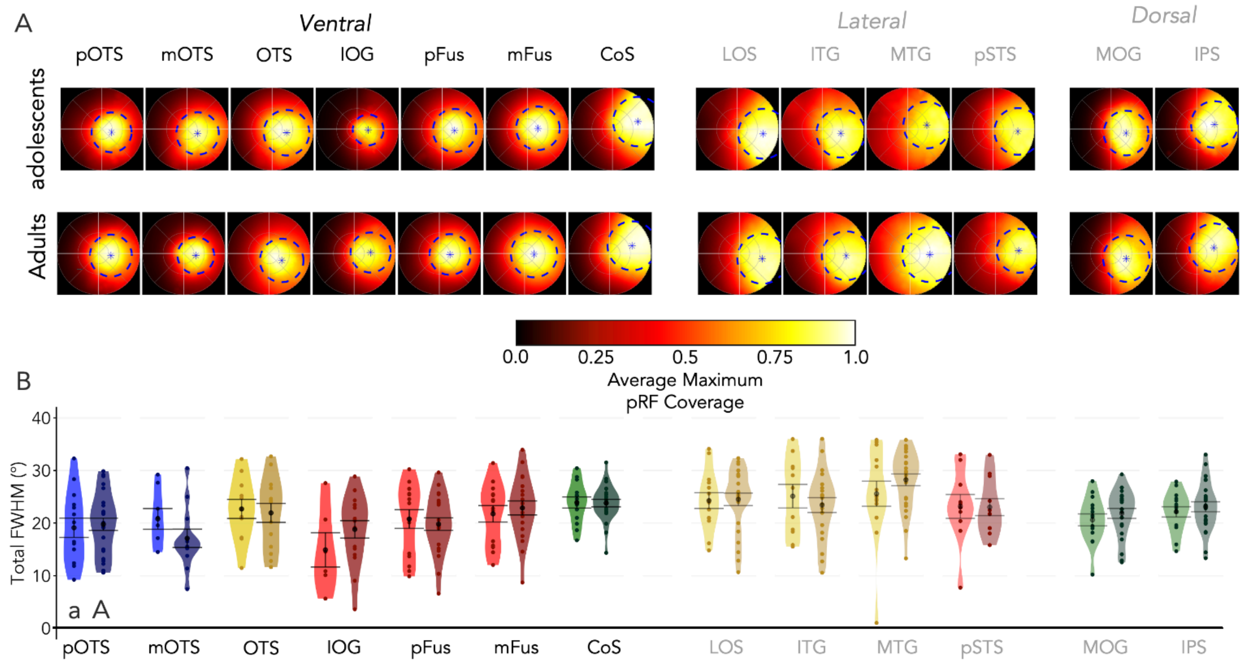

**Supplementary Figure 9. Visual field coverage (VFC) of left category-selective ROIs** (A) Average VFC of each ROI in the left hemisphere across adolescents (top row) and adults (bottom row) with a warm gradient ranging from higher coverage in white and lower coverage in black. VFC is calculated as the maximum response of pRFs covering each point in the visual field for each participant and is then averaged across participants in the group. *Blue asterisks:* average center of mass of the VFC. *Blue dotted lines:* average full-width half max (FWHM) of the coverage. (B) Violin plots of total FWHM in visual degrees for category-selective ROIs in the left hemisphere ventral, lateral, and dorsal streams in adolescents (a; lighter colors ) and adults (A; darker colors. *Black circle*: mean. *Error bars:* ± SE

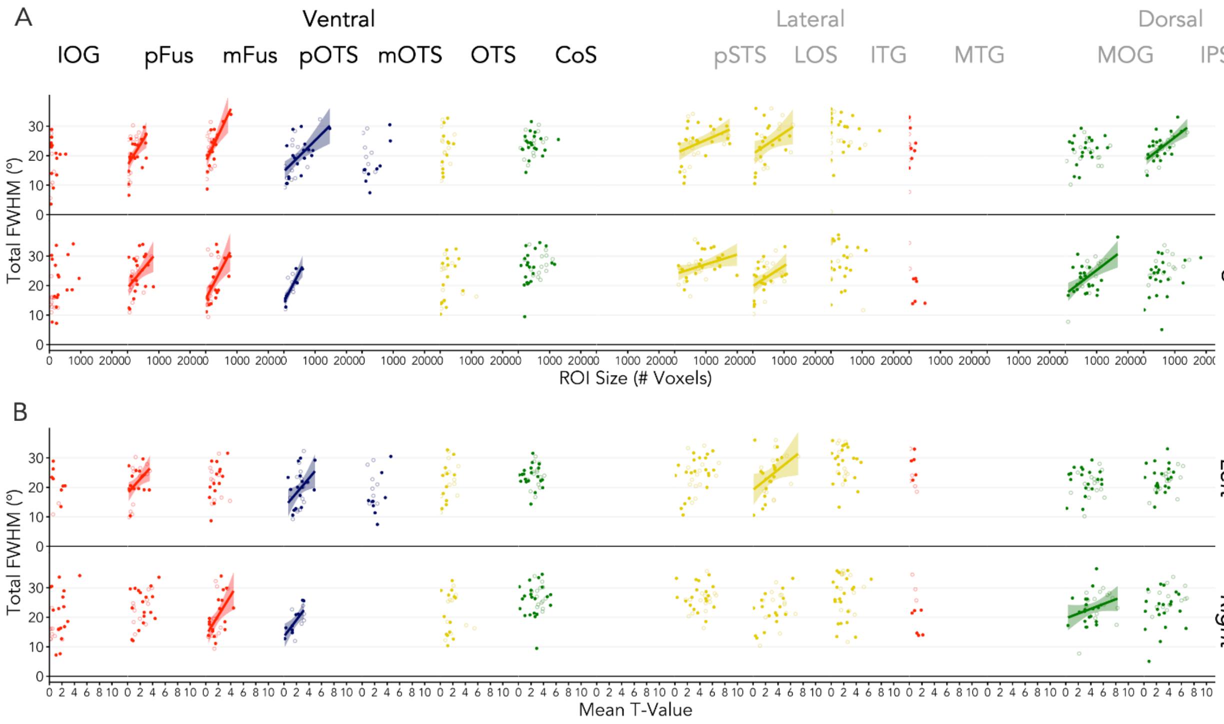

**Supplementary Figure 10. Relationship between Total FWHM and Category Selectivity.** A. Linear relationships between total FWHM and ROI size for each category-selective ROI. Each dot is a participant; adolescents are colored in lighter colors. Lines indicate a significant relationship between total FWHM and ROI size. B. Same as A but for mean t-value.

### Tables

**Supplementary Table 1. Differences in pRF parameters across early visual cortex.** Results of LMMs (*parameter ~ Age Group x ROI (V1/V2/V3) x Hemisphere + (1| participant)*) examining differences across factors: age group, ROI, and hemisphere in early visual cortex (V1, V2, V3) for pRF center x-position, pRF center y-position, pRF eccentricity, and pRF size. We find no significant effect of age group across all parameters in early visual cortex.

| Analysis | model term | df1 | df2 | F.ratio | p.value |
|---|---|---|---|---|---|
| LMM: pRF centers X-position (Age Group) | group | 1 | 40 | 0.009 | 0.9241464 |
| | ROI | 2 | 200 | 0.264 | 0.7679433 |
| | hemi | 1 | 200 | 10756.457 | 7.777e-176 |
| | group:ROI | 2 | 200 | 0.374 | 0.6885144 |
| | group:hemi | 1 | 200 | 0.028 | 0.8674489 |
| | ROI:hemi | 2 | 200 | 1.202 | 0.3027704 |
| | group:ROI:hemi | 2 | 200 | 1.066 | 0.3463284 |
| LMM: pRF centers Y-position (Age Group) | group | 1 | 40 | 0.011 | 0.9168957 |
| | ROI | 2 | 200 | 0.358 | 0.6994219 |
| | hemi | 1 | 200 | 0.683 | 0.4095495 |
| | group:ROI | 2 | 200 | 2.459 | 0.08813249 |
| | group:hemi | 1 | 200 | 0.274 | 0.6013173 |
| | ROI:hemi | 2 | 200 | 1.214 | 0.2991804 |
| | group:ROI:hemi | 2 | 200 | 1.152 | 0.3181497 |
| LMM: pRF centers Eccentricity (Age Group) | group | 1 | 40 | 0.050 | 0.8234434 |
| | ROI | 2 | 200 | 0.721 | 0.4874026 |
| | hemi | 1 | 200 | 22.907 | 3.299e-06 |
| | group:ROI | 2 | 200 | 1.911 | 0.1506917 |
| | group:hemi | 1 | 200 | 0.014 | 0.9044544 |
| | ROI:hemi | 2 | 200 | 0.140 | 0.8690739 |
| | group:ROI:hemi | 2 | 200 | 0.748 | 0.4747786 |
| LMM: pRF size (Age Group) | group | 1 | 40 | 0.154 | 0.6970016 |
| | ROI | 2 | 200 | 434.138 | 1.717e-73 |
| | hemi | 1 | 200 | 10.789 | 0.001205672 |
| | group:ROI | 2 | 200 | 0.334 | 0.7166424 |
| | group:hemi | 1 | 200 | 0.038 | 0.8461736 |
| | ROI:hemi | 2 | 200 | 8.805 | 0.0002164277 |
| | group:ROI:hemi | 2 | 200 | 1.016 | 0.3637664 |

**Supplementary Table 2. Relationship between pRF size v. eccentricity.**

Results of LMM (*pRF Size ~ Eccentricity x Age Group x ROI (V1/V2/V3) x Hemisphere + (1| participant)*) examining differences in pRF size across factors: eccentricity, age group, ROI, and hemisphere in early visual cortex (V1, V2, V3).

| Analysis | model term | df1 | df2 | F.ratio | p.value |
|---|---|---|---|---|---|
| LMM: Size v. Ecc Slopes | group | 1 | 33 | 0.018 | 0.8933011 |
| | ROI | 2 | 165 | 79.216 | 7.681e-25 |
| | hemi | 1 | 165 | 0.083 | 0.7733338 |
| | group:ROI | 2 | 165 | 0.694 | 0.5009582 |
| | group:hemi | 1 | 165 | 0.145 | 0.7034837 |
| | ROI:hemi | 2 | 165 | 2.689 | 0.07090125 |
| | group:ROI:hemi | 2 | 165 | 1.701 | 0.1856676 |
| LMM: Size v. Ecc Intercepts | group | 1 | 33 | 0.291 | 0.5930899 |
| | ROI | 2 | 165 | 158.376 | 4.069e-39 |
| | hemi | 1 | 165 | 11.113 | 0.001058557 |
| | group:ROI | 2 | 165 | 0.983 | 0.3764351 |
| | group:hemi | 1 | 165 | 0.055 | 0.8150063 |
| | ROI:hemi | 2 | 165 | 2.651 | 0.07358527 |
| | group:ROI:hemi | 2 | 165 | 0.267 | 0.7663083 |

**Supplementary Table 3. Participant counts for ROIs across hemispheres and age groups.** Number of adolescents (out of 15) and adults (out of 27) with left and right hemisphere ventral (top), lateral (middle), and dorsal (bottom) ROIs that had ten or more voxels modulated by the Toonotopy experiment. Rows with asterisks (*) denote regions derived from maximal probability maps.

| | Left Hemisphere | | Right Hemisphere | |
|---|---|---|---|---|
| ROI | adolescents | Adults | adolescents | Adults |
| IOG-faces | 6 | 16 | 11 | 20 |
| pFus-faces | 15 | 22 | 12 | 24 |
| mFus-faces | 14 | 20 | 12 | 21 |
| pOTS-words | 14 | 26 | 6 | 15 |
| mOTS-words | 8 | 12 | | |
| OTS-bodies | 11 | 14 | 11 | 19 |
| CoS-places | 14 | 24 | 14 | 25 |
| LOS-bodies | 14 | 25 | 15 | 24 |
| ITG-bodies | 11 | 21 | 13 | 23 |
| MTG-bodies* | 14 | 25 | 14 | 27 |
| pSTS-faces* | 10 | 11 | 4 | 9 |
| MOG-places | 15 | 23 | 15 | 25 |
| IPS-places | 15 | 25 | 15 | 25 |

**Supplementary Table 4. Differences in proportion of voxels with greater than 20% variance explained by the Toonotopy experiment.**

*Top:* Results of LMM (*proportion voxels with VE > 20% ~ Stream x Category x Age Group x Hemisphere +(1|participant)*) examining differences in proportion of voxels with >20% variance explained across factors: age group, stream, category, and hemisphere in ventral (excluding word-selective ROIs) and dorsal-lateral ROIs. *Bottom:* Results of ventral LMM (*proportion voxels with VE > 20% ~ Stream x Category x Age Group x Hemisphere +(1|participant)*) examining differences in proportion of voxels with >20% variance explained across factors: age group, category, and hemisphere in ventral ROIs.

| Analysis | model term | df1 | df2 | F.ratio | p.value |
|---|---|---|---|---|---|
| LMM: Proportion Voxels with Variance Explained > 20% | group | 1 | 56.26 | 3.078 | 0.08480231 |
| | stream | 1 | 690.55 | 13.232 | 0.0002956951 |
| | category | 2 | 684.58 | 31.455 | 8.527e-14 |
| | hemi | 1 | 689.67 | 14.153 | 0.0001828055 |
| | group:stream | 1 | 690.55 | 0.428 | 0.5131789 |
| | group:category | 2 | 684.58 | 4.638 | 0.009982825 |
| | group:hemi | 1 | 689.67 | 0.009 | 0.9232085 |
| | stream:category | 2 | 683.90 | 72.186 | 3.603e-29 |
| | stream:hemi | 1 | 689.21 | 0.662 | 0.4162766 |
| | category:hemi | 2 | 683.51 | 0.302 | 0.7391547 |
| | group:stream:category | 2 | 683.90 | 2.493 | 0.08337202 |
| | group:stream:hemi | 1 | 689.21 | 0.335 | 0.563069 |
| | group:category:hemi | 2 | 683.51 | 0.190 | 0.826909 |
| | stream:category:hemi | 2 | 682.32 | 4.832 | 0.008245274 |
| | group:stream:category:hemi | 2 | 682.32 | 1.943 | 0.1440752 |
| Ventral LMM: Proportion Voxels with Variance Explained > 20% | group | 1 | 47.13 | 2.753 | 0.1037378 |
| | category | 3 | 355.87 | 36.750 | 1.034e-20 |
| | hemi | 1 | 359.34 | 9.386 | 0.002351698 |
| | group:category | 3 | 355.87 | 0.854 | 0.4651188 |
| | group:hemi | 1 | 359.34 | 1.070 | 0.301635 |
| | category:hemi | 3 | 354.93 | 3.754 | 0.01119117 |
| | group:category:hemi | 3 | 354.93 | 0.879 | 0.452 |

**Supplementary Table 5. Results of Bootstrap KS-tests.**
Results of bootstrapped (n = 10000) KS-tests testing for differences in pRF x-position, y-position, eccentricity, and size between adolescents and adults. Asterisks: significant difference between groups after Bonferroni correction for multiple comparisons.

| | Left hemisphere | | | Right hemisphere | | |
|---|---|---|---|---|---|---|
| | Mean KS (D) | 95% CI | Bootstrap p | Mean KS (D) | 95% CI | Bootstrap p |
| X | | | | | | |
| pOTS - words | 0.081 | (0.05, 0.13) | 9.79e-01 | 0.149 | (0.08, 0.25) | 9.73e-01 |
| mOTS - words | 0.323 | (0.18, 0.46) | 3.21e-01 | 0.335 | (0.20, 0.50) | 5.98e-01 |
| OTS - bodies | 0.189 | (0.12, 0.31) | 9.88e-01 | 0.141 | (0.08, 0.23) | 9.96e-01 |
| IOG - faces | 0.218 | (0.11, 0.35) | 9.35e-01 | 0.125 | (0.07, 0.21) | 9.48e-01 |
| pFus - faces | 0.314 | (0.22, 0.41) | 7.35e-03 ✱ | 0.111 | (0.06, 0.17) | 9.60e-01 |
| mFus - faces | 0.154 | (0.08, 0.25) | 9.16e-01 | 0.127 | (0.07, 0.21) | 9.55e-01 |
| CoS - places | 0.279 | (0.20, 0.36) | 7.15e-03 ✱ | 0.107 | (0.06, 0.17) | 8.32e-01 |
| LOS - limbs | 0.142 | (0.08, 0.21) | 4.02e-01 | 0.086 | (0.04, 0.14) | 8.62e-01 |
| ITG - limbs | 0.110 | (0.06, 0.17) | 9.54e-01 | 0.085 | (0.04, 0.15) | 9.80e-01 |
| MTG - limbs | 0.107 | (0.06, 0.18) | 9.72e-01 | 0.131 | (0.08, 0.19) | 7.71e-01 |
| pSTS - faces | 0.200 | (0.10, 0.34) | 9.80e-01 | 0.183 | (0.10, 0.32) | 9.93e-01 |
| MOG - places | 0.066 | (0.04, 0.11) | 9.76e-01 | 0.077 | (0.05, 0.12) | 9.04e-01 |
| IPS - places | 0.159 | (0.11, 0.22) | 1.30e-01 | 0.172 | (0.12, 0.23) | 1.35e-02 ✱ |
| Y | | | | | | |
| pOTS - words | 0.073 | (0.04, 0.12) | 9.85e-01 | 0.137 | (0.07, 0.23) | 9.83e-01 |
| mOTS - words | 0.329 | (0.21, 0.45) | 2.69e-01 | 0.560 | (0.40, 0.70) | 5.35e-03 ✱ |
| OTS - bodies | 0.272 | (0.12, 0.44) | 7.78e-01 | 0.158 | (0.09, 0.26) | 9.69e-01 |
| IOG - faces | 0.209 | (0.11, 0.32) | 9.61e-01 | 0.238 | (0.15, 0.33) | 1.49e-01 |
| pFus - faces | 0.134 | (0.06, 0.22) | 9.19e-01 | 0.106 | (0.05, 0.17) | 9.66e-01 |
| mFus - faces | 0.447 | (0.34, 0.56) | 5.00e-05 ✱ | 0.282 | (0.19, 0.38) | 4.44e-02 ✱ |
| CoS - places | 0.124 | (0.06, 0.21) | 8.86e-01 | 0.103 | (0.05, 0.17) | 8.63e-01 |
| LOS - limbs | 0.138 | (0.07, 0.21) | 4.46e-01 | 0.086 | (0.04, 0.14) | 8.37e-01 |
| ITG - limbs | 0.179 | (0.11, 0.26) | 3.39e-01 | 0.183 | (0.11, 0.26) | 2.27e-01 |
| MTG - limbs | 0.166 | (0.10, 0.24) | 5.97e-01 | 0.195 | (0.12, 0.27) | 1.25e-01 |
| pSTS - faces | 0.375 | (0.20, 0.54) | 3.93e-01 | 0.201 | (0.12, 0.32) | 9.91e-01 |
| MOG - places | 0.088 | (0.04, 0.14) | 8.10e-01 | 0.134 | (0.09, 0.18) | 1.01e-01 |
| IPS - places | 0.074 | (0.04, 0.12) | 9.89e-01 | 0.081 | (0.05, 0.12) | 9.33e-01 |
| eccen | | | | | | |
| pOTS - words | 0.082 | (0.05, 0.12) | 9.85e-01 | 0.153 | (0.09, 0.24) | 9.77e-01 |
| mOTS - words | 0.310 | (0.17, 0.45) | 3.94e-01 | 0.235 | (0.14, 0.36) | 9.70e-01 |
| OTS - bodies | 0.186 | (0.10, 0.31) | 9.90e-01 | 0.176 | (0.09, 0.30) | 9.08e-01 |
| IOG - faces | 0.232 | (0.13, 0.37) | 8.85e-01 | 0.145 | (0.09, 0.21) | 9.25e-01 |
| pFus - faces | 0.283 | (0.18, 0.38) | 4.43e-02 ✱ | 0.089 | (0.05, 0.15) | 9.91e-01 |
| mFus - faces | 0.132 | (0.08, 0.21) | 9.84e-01 | 0.116 | (0.06, 0.21) | 9.66e-01 |
| CoS - places | 0.282 | (0.20, 0.37) | 5.95e-03 ✱ | 0.103 | (0.05, 0.16) | 8.71e-01 |
| LOS - limbs | 0.161 | (0.10, 0.22) | 1.76e-01 | 0.111 | (0.06, 0.16) | 5.62e-01 |
| ITG - limbs | 0.117 | (0.06, 0.19) | 8.93e-01 | 0.125 | (0.07, 0.20) | 8.07e-01 |
| MTG - limbs | 0.133 | (0.07, 0.20) | 8.66e-01 | 0.146 | (0.08, 0.22) | 5.87e-01 |
| pSTS - faces | 0.174 | (0.10, 0.30) | 9.96e-01 | 0.232 | (0.12, 0.36) | 9.62e-01 |
| MOG - places | 0.068 | (0.04, 0.11) | 9.67e-01 | 0.094 | (0.05, 0.14) | 6.69e-01 |
| IPS - places | 0.128 | (0.08, 0.18) | 5.20e-01 | 0.087 | (0.04, 0.14) | 8.27e-01 |
| size | | | | | | |
| pOTS - words | 0.246 | (0.18, 0.31) | 3.50e-04 ✱ | 0.237 | (0.12, 0.36) | 5.47e-01 |
| mOTS - words | 0.235 | (0.14, 0.35) | 8.35e-01 | 0.194 | (0.10, 0.32) | 9.87e-01 |
| OTS - bodies | 0.199 | (0.10, 0.33) | 9.81e-01 | 0.199 | (0.09, 0.33) | 8.29e-01 |
| IOG - faces | 0.213 | (0.10, 0.35) | 9.29e-01 | 0.198 | (0.12, 0.28) | 4.56e-01 |
| pFus - faces | 0.249 | (0.15, 0.35) | 1.41e-01 | 0.130 | (0.07, 0.21) | 8.47e-01 |
| mFus - faces | 0.220 | (0.12, 0.32) | 5.23e-01 | 0.338 | (0.24, 0.43) | 8.50e-04 ✱ |
| CoS - places | 0.296 | (0.21, 0.38) | 2.55e-03 ✱ | 0.133 | (0.07, 0.20) | 5.63e-01 |
| LOS - limbs | 0.092 | (0.06, 0.14) | 9.62e-01 | 0.091 | (0.04, 0.14) | 8.05e-01 |
| ITG - limbs | 0.085 | (0.04, 0.14) | 9.92e-01 | 0.151 | (0.09, 0.22) | 5.33e-01 |
| MTG - limbs | 0.156 | (0.08, 0.24) | 6.73e-01 | 0.132 | (0.09, 0.19) | 7.87e-01 |
| pSTS - faces | 0.172 | (0.10, 0.30) | 9.96e-01 | 0.443 | (0.30, 0.60) | 1.12e-01 |
| MOG - places | 0.059 | (0.03, 0.10) | 9.91e-01 | 0.112 | (0.06, 0.17) | 3.67e-01 |
| IPS - places | 0.222 | (0.15, 0.29) | 2.75e-03 ✱ | 0.205 | (0.15, 0.26) | 2.50e-04 ✱ |

✱ Bonferroni-corrected significance (bootstrap *p* < stream threshold)
Stream thresholds: Ventral = 0.0038; Lateral/Dorsal = 0.0042

**Supplementary Table 6. Differences in pRF y-position.**

*Analysis 1:* Results of LMM (*pRF y-position ~ Stream x Category x Age Group x Hemisphere +(1|participant)*) examining differences in pRF y-position explained across factors: age group, stream, category, and hemisphere in ventral (excluding word-selective ROIs) and dorsal-lateral ROIs. *Analysis 2:* Results of ventral LMM (*pRF y-position ~ Stream x Category x Age Group x Hemisphere +(1|participant)*) examining differences in pRF y-position explained across factors: age group, category, and hemisphere in ventral ROIs. *Analysis 3:* Same as Analysis 1 but with age as a continuous variable. *Analysis 4:* Same as Analysis 2 but with age as a continuous variable.

| Analysis | model term | df1 | df2 | F.ratio | p.value |
|---|---|---|---|---|---|
| LMM: pRF centers Y-position (Age Group) | group | 1 | 49.27 | 1.097 | 0.300016 |
| | stream | 1 | 686.75 | 3.333 | 0.06834519 |
| | category | 2 | 681.25 | 54.018 | 1.677e-22 |
| | hemi | 1 | 685.54 | 28.482 | 1.287e-07 |
| | group:stream | 1 | 686.75 | 1.561 | 0.2119513 |
| | group:category | 2 | 681.25 | 1.130 | 0.3237693 |
| | group:hemi | 1 | 685.54 | 0.033 | 0.8553255 |
| | stream:category | 2 | 680.53 | 12.150 | 6.537e-06 |
| | stream:hemi | 1 | 685.24 | 0.431 | 0.5115772 |
| | category:hemi | 2 | 680.10 | 3.440 | 0.03263797 |
| | group:stream:category | 2 | 680.53 | 0.633 | 0.5315631 |
| | group:stream:hemi | 1 | 685.24 | 0.154 | 0.6950682 |
| | group:category:hemi | 2 | 680.10 | 0.117 | 0.8898049 |
| | stream:category:hemi | 2 | 679.48 | 2.159 | 0.1162758 |
| | group:stream:category:hemi | 2 | 679.48 | 0.417 | 0.6593742 |
| Ventral LMM: pRF centers Y-position (Age Group) | group | 1 | 46.06 | 0.277 | 0.6009889 |
| | category | 3 | 353.99 | 54.104 | 8.256e-29 |
| | hemi | 1 | 357.51 | 50.868 | 5.518e-12 |
| | group:category | 3 | 353.99 | 0.156 | 0.9260235 |
| | group:hemi | 1 | 357.51 | 0.059 | 0.8077871 |
| | category:hemi | 3 | 353.12 | 3.331 | 0.01973055 |
| | group:category:hemi | 3 | 353.12 | 0.262 | 0.8526528 |
| LMM: pRF centers Y-position (Age Continuous) | age | 1 | 50.43 | 0.948 | 0.3349186 |
| | stream | 1 | 688.15 | 2.371 | 0.1240674 |
| | category | 2 | 681.65 | 58.259 | 4.386e-24 |
| | hemi | 1 | 686.71 | 29.892 | 6.4e-08 |
| | age:stream | 1 | 688.27 | 3.409 | 0.06527422 |
| | age:category | 2 | 681.53 | 0.742 | 0.4763231 |
| | age:hemi | 1 | 684.62 | 0.050 | 0.8227047 |
| | stream:category | 2 | 681.21 | 13.988 | 1.113e-06 |
| | stream:hemi | 1 | 685.11 | 0.324 | 0.5693828 |
| | category:hemi | 2 | 680.49 | 3.291 | 0.03779714 |
| | age:stream:category | 2 | 680.57 | 0.783 | 0.4572741 |
| | age:stream:hemi | 1 | 686.66 | 0.158 | 0.6915707 |
| | age:category:hemi | 2 | 680.14 | 0.325 | 0.7223864 |
| | stream:category:hemi | 2 | 679.78 | 1.810 | 0.1644058 |
| | age:stream:category:hemi | 2 | 679.82 | 1.026 | 0.3589187 |
| Ventral LMM: pRF centers Y-position (Age Continuous) | age | 1 | 48.12 | 0.007 | 0.9349859 |
| | category | 3 | 354.31 | 56.999 | 4.342e-30 |
| | hemi | 1 | 355.72 | 53.864 | 1.472e-12 |
| | age:category | 3 | 354.07 | 0.064 | 0.9789812 |
| | age:hemi | 1 | 359.80 | 0.079 | 0.7790408 |
| | category:hemi | 3 | 353.16 | 3.785 | 0.01073501 |
| | age:category:hemi | 3 | 353.29 | 0.505 | 0.6790506 |

**Supplementary Table 7. Differences in pRF eccentricity.**
*Analysis 1:* Results of LMM (*Eccentricity ~ Stream x Category x Age Group x Hemisphere +(1|participant)*) examining differences in pRF eccentricity explained across factors: age group, stream, category, and hemisphere in ventral (excluding word-selective ROIs) and dorsal-lateral ROIs. *Analysis 2:* Results of ventral LMM (*Eccentricity ~ Stream x Category x Age Group x Hemisphere +(1|participant)*) examining differences in pRF eccentricity explained across factors: age group, category, and hemisphere in ventral ROIs. *Analysis 3:* Same as Analysis 1 but with age as a continuous variable. *Analysis 4:* Same as Analysis 2 but with age as a continuous variable.

| Analysis | model term | df1 | df2 | F.ratio | p.value |
|---|---|---|---|---|---|
| LMM: pRF centers Eccentricity (Age Group) | group | 1 | 56.38 | 0.387 | 0.5364263 |
| | stream | 1 | 690.80 | 84.705 | 4.052e-19 |
| | category | 2 | 682.44 | 6.219 | 0.002106673 |
| | hemi | 1 | 689.28 | 58.661 | 6.359e-14 |
| | group:stream | 1 | 690.80 | 0.107 | 0.7440599 |
| | group:category | 2 | 682.44 | 1.270 | 0.2814686 |
| | group:hemi | 1 | 689.28 | 0.089 | 0.7656275 |
| | stream:category | 2 | 681.10 | 106.563 | 5.427e-41 |
| | stream:hemi | 1 | 688.93 | 1.428 | 0.2325535 |
| | category:hemi | 2 | 680.54 | 7.585 | 0.0005523304 |
| | group:stream:category | 2 | 681.10 | 0.330 | 0.7190713 |
| | group:stream:hemi | 1 | 688.93 | 0.477 | 0.4900051 |
| | group:category:hemi | 2 | 680.54 | 0.094 | 0.9103302 |
| | stream:category:hemi | 2 | 679.44 | 10.708 | 2.639e-05 |
| | group:stream:category:hemi | 2 | 679.44 | 0.228 | 0.7962663 |
| Ventral LMM: pRF centers Eccentricity (Age Group) | group | 1 | 52.03 | 0.372 | 0.5443443 |
| | category | 3 | 356.94 | 84.643 | 2.206e-41 |
| | hemi | 1 | 362.92 | 21.314 | 5.42e-06 |
| | group:category | 3 | 356.94 | 1.186 | 0.3148169 |
| | group:hemi | 1 | 362.92 | 0.004 | 0.9522864 |
| | category:hemi | 3 | 355.07 | 5.074 | 0.001880144 |
| | group:category:hemi | 3 | 355.07 | 0.447 | 0.7192879 |
| LMM: pRF centers Eccentricity (Age Continuous) | age | 1 | 57.93 | 0.716 | 0.4008128 |
| | stream | 1 | 692.35 | 90.575 | 2.894e-20 |
| | category | 2 | 682.85 | 5.285 | 0.005275028 |
| | hemi | 1 | 690.69 | 64.132 | 4.943e-15 |
| | age:stream | 1 | 693.07 | 0.748 | 0.3875659 |
| | age:category | 2 | 682.73 | 1.893 | 0.1514428 |
| | age:hemi | 1 | 687.54 | 0.094 | 0.7590919 |
| | stream:category | 2 | 681.96 | 110.021 | 3.877e-42 |
| | stream:hemi | 1 | 688.01 | 2.158 | 0.1423128 |
| | category:hemi | 2 | 681.05 | 8.350 | 0.0002613315 |
| | age:stream:category | 2 | 680.95 | 0.187 | 0.8296404 |
| | age:stream:hemi | 1 | 691.29 | 0.513 | 0.4742924 |
| | age:category:hemi | 2 | 680.35 | 0.006 | 0.9936298 |
| | stream:category:hemi | 2 | 679.79 | 10.822 | 2.363e-05 |
| | age:stream:category:hemi | 2 | 679.79 | 0.103 | 0.9022713 |
| Ventral LMM: pRF centers Eccentricity (Age Continuous) | age | 1 | 55.50 | 0.758 | 0.3877128 |
| | category | 3 | 357.98 | 85.853 | 7.218e-42 |
| | hemi | 1 | 360.48 | 23.931 | 1.508e-06 |
| | age:category | 3 | 357.33 | 1.364 | 0.2536275 |
| | age:hemi | 1 | 367.18 | 0.397 | 0.5290587 |
| | category:hemi | 3 | 355.82 | 4.879 | 0.002448944 |
| | age:category:hemi | 3 | 355.51 | 0.107 | 0.9562231 |

**Supplementary Table 8. Differences in pRF size.**
*Analysis 1:* Results of LMM (*pRF size ~ Stream x Category x Age Group x Hemisphere +(1|participant)*) examining differences in pRF size explained across factors: age group, stream, category, and hemisphere in ventral (excluding word-selective ROIs) and dorsal-lateral ROIs. *Analysis 2:* Results of ventral LMM (*pRF size ~ Stream x Category x Age Group x Hemisphere +(1|participant)*) examining differences in pRF size explained across factors: age group, category, and hemisphere in ventral ROIs. *Analysis 3:* Same as Analysis 1 but with age as a continuous variable. *Analysis 4:* Same as Analysis 2 but with age as a continuous variable.

| Analysis | model term | df1 | df2 | F.ratio | p.value |
|---|---|---|---|---|---|
| LMM: pRF Size (Age Group) | group | 1 | 66.98 | 5.627 | 0.02057086 |
| | stream | 1 | 695.56 | 35.679 | 3.713e-09 |
| | category | 2 | 687.64 | 21.703 | 7.242e-10 |
| | hemi | 1 | 694.18 | 1.709 | 0.1915752 |
| | group:stream | 1 | 695.56 | 3.291 | 0.07008061 |
| | group:category | 2 | 687.64 | 1.230 | 0.2930589 |
| | group:hemi | 1 | 694.18 | 9.838 | 0.001781559 |
| | stream:category | 2 | 686.30 | 25.697 | 1.731e-11 |
| | stream:hemi | 1 | 693.88 | 0.862 | 0.3533844 |
| | category:hemi | 2 | 685.79 | 0.530 | 0.5889053 |
| | group:stream:category | 2 | 686.30 | 1.480 | 0.2284042 |
| | group:stream:hemi | 1 | 693.88 | 4.064 | 0.04418602 |
| | group:category:hemi | 2 | 685.79 | 2.951 | 0.05296232 |
| | stream:category:hemi | 2 | 684.71 | 0.575 | 0.5629769 |
| | group:stream:category:hemi | 2 | 684.71 | 3.620 | 0.02730318 |
| Ventral LMM: pRF Size (Age Group) | group | 1 | 59.23 | 3.952 | 0.05144717 |
| | category | 3 | 361.00 | 19.098 | 1.622e-11 |
| | hemi | 1 | 366.63 | 1.316 | 0.2520593 |
| | group:category | 3 | 361.00 | 0.636 | 0.5921668 |
| | group:hemi | 1 | 366.63 | 2.508 | 0.1141252 |
| | category:hemi | 3 | 359.17 | 0.342 | 0.7948369 |
| | group:category:hemi | 3 | 359.17 | 1.576 | 0.1948032 |
| LMM: pRF Size (Age Continuous) | age | 1 | 68.39 | 4.237 | 0.0433552 |
| | stream | 1 | 696.73 | 33.338 | 1.167e-08 |
| | category | 2 | 688.04 | 20.460 | 2.337e-09 |
| | hemi | 1 | 695.28 | 0.511 | 0.4751575 |
| | age:stream | 1 | 697.46 | 3.790 | 0.05196879 |
| | age:category | 2 | 687.94 | 1.063 | 0.3461186 |
| | age:hemi | 1 | 692.40 | 7.377 | 0.006771967 |
| | stream:category | 2 | 687.19 | 24.930 | 3.531e-11 |
| | stream:hemi | 1 | 692.80 | 0.366 | 0.5451499 |
| | category:hemi | 2 | 686.37 | 0.403 | 0.6683421 |
| | age:stream:category | 2 | 686.25 | 1.736 | 0.1770626 |
| | age:stream:hemi | 1 | 695.94 | 3.931 | 0.04779728 |
| | age:category:hemi | 2 | 685.69 | 2.540 | 0.07958764 |
| | stream:category:hemi | 2 | 685.18 | 0.208 | 0.8123016 |
| | age:stream:category:hemi | 2 | 685.17 | 3.316 | 0.03689149 |
| Ventral LMM: pRF Size (Age Continuous) | age | 1 | 62.68 | 1.473 | 0.229466 |
| | category | 3 | 361.81 | 18.165 | 5.317e-11 |
| | hemi | 1 | 364.14 | 0.507 | 0.4769429 |
| | age:category | 3 | 361.18 | 0.146 | 0.9322512 |
| | age:hemi | 1 | 370.31 | 0.818 | 0.3663343 |
| | category:hemi | 3 | 359.78 | 0.462 | 0.709319 |
| | age:category:hemi | 3 | 359.43 | 1.194 | 0.3118894 |

**Supplementary Table 9. Differences in VFC FWHM.**

*Analysis 1:* Results of LMM (*FWHM ~ Stream x Category x Age Group x Hemisphere +(1|participant)*) examining differences in VFC FWHM explained across factors: age group, stream, category, and hemisphere in ventral (excluding word-selective ROIs) and dorsal-lateral ROIs. *Analysis 2:* Results of ventral LMM (*FWHM ~ Stream x Category x Age Group x Hemisphere +(1|participant)*) examining differences in VFC FWHM explained across factors: age group, category, and hemisphere in ventral ROIs. *Analysis 3:* Same as Analysis 1 but with age as a continuous variable. *Analysis 4:* Same as Analysis 2 but with age as a continuous variable.

| Analysis | model term | df1 | df2 | F.ratio | p.value |
|---|---|---|---|---|---|
| LMM: Total FWHM (Age Group) | group | 1 | 65.46 | 1.837 | 0.1799715 |
| | stream | 1 | 694.72 | 7.024 | 0.00822665 |
| | category | 2 | 686.50 | 4.094 | 0.01707638 |
| | hemi | 1 | 693.82 | 3.636 | 0.05695543 |
| | group:stream | 1 | 694.72 | 4.782 | 0.02909658 |
| | group:category | 2 | 686.50 | 0.883 | 0.4139856 |
| | group:hemi | 1 | 693.82 | 3.248 | 0.0719624 |
| | stream:category | 2 | 685.38 | 14.525 | 6.638e-07 |
| | stream:hemi | 1 | 693.29 | 0.227 | 0.6337889 |
| | category:hemi | 2 | 684.95 | 2.181 | 0.113698 |
| | group:stream:category | 2 | 685.38 | 5.996 | 0.002620842 |
| | group:stream:hemi | 1 | 693.29 | 7.148 | 0.007680015 |
| | group:category:hemi | 2 | 684.95 | 0.859 | 0.4242138 |
| | stream:category:hemi | 2 | 683.10 | 0.624 | 0.5359071 |
| | group:stream:category:hemi | 2 | 683.10 | 3.513 | 0.03034913 |
| Ventral LMM: Total FWHM (Age Group) | group | 1 | 52.03 | 0.013 | 0.9103476 |
| | category | 3 | 358.41 | 13.639 | 1.914e-08 |
| | hemi | 1 | 363.33 | 0.781 | 0.3772795 |
| | group:category | 3 | 358.41 | 1.536 | 0.2049264 |
| | group:hemi | 1 | 363.33 | 0.338 | 0.5614568 |
| | category:hemi | 3 | 356.91 | 1.327 | 0.2654512 |
| | group:category:hemi | 3 | 356.91 | 0.767 | 0.5131287 |
| LMM: Total FWHM (Age Continuous) | group | 1 | 65.46 | 1.837 | 0.1799715 |
| | stream | 1 | 694.72 | 7.024 | 0.00822665 |
| | category | 2 | 686.50 | 4.094 | 0.01707638 |
| | hemi | 1 | 693.82 | 3.636 | 0.05695543 |
| | group:stream | 1 | 694.72 | 4.782 | 0.02909658 |
| | group:category | 2 | 686.50 | 0.883 | 0.4139856 |
| | group:hemi | 1 | 693.82 | 3.248 | 0.0719624 |
| | stream:category | 2 | 685.38 | 14.525 | 6.638e-07 |
| | stream:hemi | 1 | 693.29 | 0.227 | 0.6337889 |
| | category:hemi | 2 | 684.95 | 2.181 | 0.113698 |
| | group:stream:category | 2 | 685.38 | 5.996 | 0.002620842 |
| | group:stream:hemi | 1 | 693.29 | 7.148 | 0.007680015 |
| | group:category:hemi | 2 | 684.95 | 0.859 | 0.4242138 |
| | stream:category:hemi | 2 | 683.10 | 0.624 | 0.5359071 |
| | group:stream:category:hemi | 2 | 683.10 | 3.513 | 0.03034913 |
| Ventral LMM: Total FWHM (Age Continuous) | group | 1 | 52.03 | 0.013 | 0.9103476 |
| | category | 3 | 358.41 | 13.639 | 1.914e-08 |
| | hemi | 1 | 363.33 | 0.781 | 0.3772795 |
| | group:category | 3 | 358.41 | 1.536 | 0.2049264 |
| | group:hemi | 1 | 363.33 | 0.338 | 0.5614568 |
| | category:hemi | 3 | 356.91 | 1.327 | 0.2654512 |
| | group:category:hemi | 3 | 356.91 | 0.767 | 0.5131287 |

**Supplementary Table 10. Differences in category selectivity – ROI Size.**
*Analysis 1:* Results of LMM (*ROI Size ~ Stream x Category x Age Group x Hemisphere +(1|participant)*) examining differences in category selectivity (ROI size measured in number of voxels) explained across factors: age group, stream, category, and hemisphere in ventral (excluding word-selective ROIs) and dorsal-lateral ROIs. *Analysis 2:* Results of ventral LMM *ROI Size ~ Category x Age Group x Hemisphere +(1|participant)*) examining differences in category selectivity (ROI Size) explained across factors: age group, category, and hemisphere in ventral ROIs. *Analysis 3:* Same as Analysis 1 but with age as a continuous variable. *Analysis 4:* Same as Analysis 2 but with age as a continuous variable.

| Analysis | model term | df1 | df2 | F.ratio | p.value |
|---|---|---|---|---|---|
| LMM: ROI Size (Age Group) | group | 1 | 45.78 | 0.012 | 0.9128806 |
| | stream | 1 | 780.50 | 9.636 | 0.001977214 |
| | category | 2 | 777.97 | 98.486 | 7.468e-39 |
| | hemi | 1 | 777.61 | 9.452 | 0.002183152 |
| | group:stream | 1 | 780.50 | 0.045 | 0.8324554 |
| | group:category | 2 | 777.97 | 5.906 | 0.00284772 |
| | group:hemi | 1 | 777.61 | 0.028 | 0.8664297 |
| | stream:category | 2 | 777.74 | 20.553 | 2.003e-09 |
| | stream:hemi | 1 | 778.21 | 1.083 | 0.2982469 |
| | category:hemi | 2 | 776.91 | 0.523 | 0.5928702 |
| | group:stream:category | 2 | 777.74 | 0.766 | 0.4653488 |
| | group:stream:hemi | 1 | 778.21 | 0.537 | 0.4640681 |
| | group:category:hemi | 2 | 776.91 | 0.621 | 0.5376555 |
| | stream:category:hemi | 2 | 777.38 | 0.237 | 0.7887317 |
| | group:stream:category:hemi | 2 | 777.38 | 0.706 | 0.4937165 |
| Ventral LMM: ROI Size (Age Group) | group | 1 | 52.89 | 1.277 | 0.2634827 |
| | category | 3 | 385.84 | 34.532 | 8.636e-20 |
| | hemi | 1 | 389.39 | 1.086 | 0.2981133 |
| | group:category | 3 | 385.84 | 4.286 | 0.005421243 |
| | group:hemi | 1 | 389.39 | 2.812 | 0.09435181 |
| | category:hemi | 3 | 384.53 | 5.975 | 0.0005462451 |
| | group:category:hemi | 3 | 384.53 | 2.221 | 0.08524093 |
| LMM: ROI Size (Age Continuous) | age | 1 | 46.62 | 0.007 | 0.9358678 |
| | stream | 1 | 781.30 | 9.848 | 0.0017637 |
| | category | 2 | 778.30 | 91.895 | 1.484e-36 |
| | hemi | 1 | 777.28 | 9.659 | 0.001952582 |
| | age:stream | 1 | 782.85 | 0.062 | 0.8029298 |
| | age:category | 2 | 777.93 | 3.163 | 0.04285377 |
| | age:hemi | 1 | 777.86 | 0.060 | 0.8067235 |
| | stream:category | 2 | 777.90 | 23.759 | 9.651e-11 |
| | stream:hemi | 1 | 777.93 | 0.754 | 0.3853995 |
| | category:hemi | 2 | 777.26 | 0.269 | 0.7643951 |
| | age:stream:category | 2 | 777.73 | 0.349 | 0.7053069 |
| | age:stream:hemi | 1 | 778.91 | 0.366 | 0.5453154 |
| | age:category:hemi | 2 | 776.70 | 0.480 | 0.6191828 |
| | stream:category:hemi | 2 | 777.42 | 0.315 | 0.7296478 |
| | age:stream:category:hemi | 2 | 777.68 | 0.420 | 0.656917 |
| Ventral LMM: ROI Size (Age Continuous) | age | 1 | 55.76 | 1.565 | 0.2161444 |
| | category | 3 | 385.98 | 31.308 | 3.905e-18 |
| | hemi | 1 | 387.60 | 0.422 | 0.5161309 |
| | age:category | 3 | 385.42 | 1.879 | 0.1326651 |
| | age:hemi | 1 | 391.85 | 2.071 | 0.1509276 |
| | category:hemi | 3 | 384.49 | 7.365 | 8.242e-05 |
| | age:category:hemi | 3 | 384.21 | 1.750 | 0.1563758 |

**Supplementary Table 11. ROI Size LMM Betas**

Using the emmeans package in R, we obtained the estimated marginal means – the ROI size for each group after adjusting for all other factors in the LMMs from Table S10 – computed the pairwise contrasts between adolescents and adults within each category, producing a set of model-derived beta estimates. *Analysis 1:* Betas of LMM (*ROI Size ~ Stream x Category x Age Group x Hemisphere +(1|participant)*) examining differences in ROI size explained across factors: age group, stream, category, and hemisphere in ventral (excluding word-selective ROIs) and dorsal-lateral ROIs. *Analysis 2:* Betas of ventral LMM *ROI Size ~ Category x Age Group x Hemisphere +(1|participant)*) examining differences in ROI size in ventral ROIs. Positive betas: increases in ROI size from adolescence to adulthood. Negative betas: decreases in ROI size from adolescence to adulthood. Asterisks: significant difference between groups.

| Category | Group Contrast | β | SE | df | t | p-value | |
|---|---|---|---|---|---|---|---|
| LMM (no words) | | | | | | | |
| bodies | Adults - adolescents | 20.947 | 65.298 | 142.8 | 0.321 | 0.749 | |
| face | Adults - adolescents | 90.493 | 61.458 | 114.7 | 1.472 | 0.144 | |
| place | Adults - adolescents | -127.950 | 62.289 | 120.8 | -2.054 | 0.042 | * |
| words | Adults - adolescents | NA | NA | NA | NA | NA | NA |
| Ventral LMM | | | | | | | |
| bodies | Adults - adolescents | -0.655 | 75.163 | 290.0 | -0.009 | 0.993 | |
| face | Adults - adolescents | 129.990 | 45.695 | 70.9 | 2.845 | 0.006 | * |
| place | Adults - adolescents | -86.370 | 65.532 | 225.9 | -1.318 | 0.189 | |
| words | Adults - adolescents | 149.791 | 74.943 | 286.7 | 1.999 | 0.047 | * |

**Supplementary Table 12. Differences in category selectivity – mean t-value.**
As a complementary analysis to the analyses in Table S10, we quantified category selectivity (mean t-value) in 10mm disks ROI centered on each category ROI and tested if selectivity varied across age group, stream, category, and hemisphere (*LMM: Mean T-Value ~ Age Group x Stream x Category x Hemisphere +(1|Participant); ventral LMM: mean t-value ~ Age Group x Category x Hemisphere +(1|Participant)*). *Analysis 1:* Results of LMM (*Mean T-Value ~ Stream x Category x Age Group x Hemisphere +(1|participant)*) examining differences in category selectivity explained across factors: age group, stream, category, and hemisphere in ventral (excluding word-selective ROIs) and dorsal-lateral ROIs. *Analysis 2:* Results of ventral LMM ((*Mean T-Value ~ Stream x Category x Age Group x Hemisphere +(1|participant)*) examining differences in category selectivity explained across factors: age group, category, and hemisphere in ventral ROIs. *Analysis 3:* Same as Analysis 1 but with age as a continuous variable. *Analysis 4:* Same as Analysis 2 but with age as a continuous variable. Like with the mean t-values, we found a significant group by category effect, indicating that the size of ROIs differentially develops across visual categories.

| Analysis | model term | df1 | df2 | F.ratio | p.value |
|---|---|---|---|---|---|
| LMM: Category Selectivity - Mean T-Value (Age Group) | group | 1 | 41.83 | 3.649 | 0.06296847 |
| | stream | 1 | 748.81 | 8.322 | 0.004028748 |
| | category | 2 | 747.33 | 151.272 | 6.884e-56 |
| | hemi | 1 | 747.13 | 16.336 | 5.853e-05 |
| | group:stream | 1 | 748.81 | 2.911 | 0.08838366 |
| | group:category | 2 | 747.33 | 15.576 | 2.359e-07 |
| | group:hemi | 1 | 747.13 | 2.156 | 0.1424783 |
| | stream:category | 2 | 747.84 | 38.637 | 1.075e-16 |
| | stream:hemi | 1 | 747.35 | 0.272 | 0.6019871 |
| | category:hemi | 2 | 747.03 | 0.623 | 0.5368672 |
| | group:stream:category | 2 | 747.84 | 1.656 | 0.1916176 |
| | group:stream:hemi | 1 | 747.35 | 0.517 | 0.4725063 |
| | group:category:hemi | 2 | 747.03 | 0.628 | 0.533842 |
| | stream:category:hemi | 2 | 747.06 | 0.283 | 0.7533162 |
| | group:stream:category:hemi | 2 | 747.06 | 0.133 | 0.8758854 |
| Ventral LMM: Category Selectivity - Mean T-Value (Age Group) | group | 1 | 44.27 | 3.380 | 0.07268811 |
| | category | 3 | 358.60 | 33.350 | 4.919e-19 |
| | hemi | 1 | 361.34 | 4.297 | 0.03889545 |
| | group:category | 3 | 358.60 | 5.101 | 0.001810918 |
| | group:hemi | 1 | 361.34 | 2.174 | 0.1412203 |
| | category:hemi | 3 | 357.30 | 1.800 | 0.1467495 |
| | group:category:hemi | 3 | 357.30 | 1.166 | 0.3226626 |
| LMM: Category Selectivity - Mean T-Value (Age Continuous) | age | 1 | 41.97 | 5.177 | 0.0280577 |
| | stream | 1 | 748.81 | 6.676 | 0.009959008 |
| | category | 2 | 747.38 | 139.277 | 3.878e-52 |
| | hemi | 1 | 746.97 | 14.457 | 0.0001550827 |
| | age:stream | 1 | 749.29 | 3.267 | 0.0710725 |
| | age:category | 2 | 747.50 | 19.315 | 6.625e-09 |
| | age:hemi | 1 | 747.30 | 3.042 | 0.08153271 |
| | stream:category | 2 | 747.69 | 38.911 | 8.392e-17 |
| | stream:hemi | 1 | 747.20 | 0.078 | 0.7795923 |
| | category:hemi | 2 | 747.12 | 1.050 | 0.3505502 |
| | age:stream:category | 2 | 747.74 | 2.944 | 0.05324479 |
| | age:stream:hemi | 1 | 747.53 | 1.374 | 0.2415394 |
| | age:category:hemi | 2 | 746.98 | 0.763 | 0.4665843 |
| | stream:category:hemi | 2 | 747.11 | 0.415 | 0.6601895 |
| | age:stream:category:hemi | 2 | 747.04 | 0.139 | 0.8702275 |
| Ventral LMM: Category Selectivity - Mean T-Value (Age Continuous) | age | 1 | 46.90 | 4.964 | 0.03071043 |
| | category | 3 | 358.58 | 29.124 | 7.107e-17 |
| | hemi | 1 | 360.17 | 3.152 | 0.07667675 |
| | age:category | 3 | 358.73 | 5.051 | 0.001939446 |
| | age:hemi | 1 | 362.97 | 1.202 | 0.2736503 |
| | category:hemi | 3 | 357.21 | 2.731 | 0.04374043 |
| | age:category:hemi | 3 | 357.08 | 0.856 | 0.4642207 |

**Supplementary Table 13. Category Selectivity versus Spatial Integration.**
*Analysis 1:* Results of LMM comparing ROI size to spatial integration (*ROI Size LMM: ROI Size ~ VFC FWHM + pRF Size + Eccentricity + Age Group + (1 | participant)*. *Analysis 2:* Results of LMM comparing mean t-value to spatial integration *(Mean T-Value LMM: Mean T-Value ~ VFC FWHM + pRF Size + Eccentricity + Age Group + (1 | participant)*

| Analysis | model term | df1 | df2 | F.ratio | p.value |
|---|---|---|---|---|---|
| LMM: ROI size ~ fwhm + size + ecc + group | fwhm | 1 | 241.05 | 52.209 | 6.535e-12 |
| | size | 1 | 237.49 | 3.278 | 0.07145696 |
| | ecc | 1 | 239.46 | 4.442 | 0.03611168 |
| | group | 1 | 38.15 | 0.493 | 0.4870366 |
| LMM: meanT ~ fwhm + size + ecc + group | fwhm | 1 | 233.13 | 24.865 | 1.201e-06 |
| | size | 1 | 235.91 | 0.083 | 0.7736299 |
| | ecc | 1 | 225.58 | 0.948 | 0.3312479 |
| | group | 1 | 37.14 | 0.038 | 0.8472222 |

**Supplementary Table 14. Category Selectivity versus Spatial Integration Betas.**
Betas for analyses in Table S13. *Analysis 1:* Betas of LMM comparing ROI size to spatial integration (*ROI Size LMM: ROI Size ~ VFC FWHM + pRF Size + Eccentricity + Age Group + (1 | participant)*. *Analysis 2:* Betas of LMM comparing mean t-value to spatial integration *(Mean T-Value LMM: Mean T-Value ~ VFC FWHM + pRF Size + Eccentricity + Age Group + (1 | participant)*

| Analysis | Term | β | SE | df | t | p | 95% CI |
|---|---|---|---|---|---|---|---|
| **LMM: ROI size ~ fwhm + size + ecc + group** | (Intercept) | -113.600 | 64.679 | 197.4 | -1.756 | 0.081 | [-241.150, 13.950] |
| | fwhm | 21.813 | 2.993 | 241.1 | 7.288 | 4.463e-12 | [ 15.918, 27.709] |
| | size | -14.526 | 7.944 | 237.9 | -1.829 | 0.069 | [ -30.175, 1.123] |
| | ecc | 12.263 | 5.785 | 239.7 | 2.120 | 0.035 | [ 0.867, 23.659] |
| | groupAdults | 26.996 | 38.411 | 41.3 | 0.703 | 0.486 | [ -50.562, 104.553] |
| **LMM: meanT ~ fwhm + size + ecc + group** | (Intercept) | -0.303 | 0.373 | 156.5 | -0.813 | 0.417 | [ -1.041, 0.434] |
| | fwhm | 0.077 | 0.015 | 233.5 | 5.009 | 1.078e-06 | [ 0.047, 0.108] |
| | size | 0.012 | 0.041 | 236.2 | 0.289 | 0.772 | [ -0.069, 0.093] |
| | ecc | 0.029 | 0.029 | 226.2 | 0.977 | 0.330 | [ -0.029, 0.086] |
| | groupAdults | -0.058 | 0.298 | 38.5 | -0.194 | 0.847 | [ -0.662, 0.546] |
